# A Comprehensive Hybrid Capture Sequencing Platform for Tick-borne Disease Detection Reveals Zoonotic Emerging Pathogens in a Weather and Vector Transition Zone

**DOI:** 10.1101/2025.06.06.658355

**Authors:** Chloe M. Murrell, Ximena A. Olarte-Castillo, Xiyu Wang, Aiman Sabaawy, Laura W. Alexander, Grace Gonzalez, Joel J. Brown, Kristina Ceres, Massimiliano S. Tagliamonte, Nagi Hashimoto, Yining Sun, Jordan D. Zehr, Gayatri Anil, Aine Lehane, Emily M. Mader, Natalie Bailey, Marie V. Lilly, Laura D. Plimpton, Ursula E. A. Bergwerff, Guillaume Reboul, Cameron Guan, Kelly L. Sams, Sita Sahithi Mullapudi, Lauren Singh, Ethan Seiz, Nisha Khatri, Braxton Sizemore, Sebastian Llanos-Soto, Rebecca Tallmadge, Patrick K. Mitchell, Renee Anderson, Ellie Bourgikos, Chantal B.F. Vogels, Alexander T. Ciota, Victoria Schnurr, Jennifer K. Grenier, Xavier Berthet, Joellen M. Lampman, Stephen G. Ladd-Wilson, Ana I. Bento, Laura C. Harrington, Maria A. Diuk-Wasser, Carla Mavian, Laura B. Goodman

## Abstract

As weather systems quickly change, vector communities and their pathogens evolve faster than assay panels can be redesigned. Additionally, PCR’s narrow target range makes it structurally unable to catch divergent or reassorted agents. We developed a hybrid capture next-generation sequencing enrichment platform that provides comprehensive detection and characterization of tick-borne agents alongside ecological vertebrate host identification with low-pass sequencing. Analytical validation demonstrated performance comparable to qPCR with superior variant tolerance and multiplexing capacity. Field deployment in subtropical, metropolitan New York detected *Anaplasma phagocytophilum* strains (n=3) linked to human granulocytic anaplasmosis and a *Babesia microti*-like species in urban raccoon (*Procyon lotor*) populations. Tick vector screening revealed a putatively novel chimeric Flavi-like virus in invasive *Haemaphysalis longicornis* ticks combining segmented and unsegmented genomic features, with codon adaptation analysis indicating strong human compatibility. Serology revealed high *Flavivirus* seropositivity in NYC raccoons, suggesting an unrecognized urban reservoir role. Blood meal analysis simultaneously revealed complex ecological pathogen transmission networks spanning multiple vertebrate hosts. This integrated surveillance system enables comprehensive pathogen discovery, real-time evolutionary monitoring, and ecological risk assessment, transforming our capacity to detect emerging tick-borne threats in rapidly changing environments and prevent spillovers.

## Main

The global distribution of arthropod vectors is changing faster than diagnostic capacity can be developed. In temperate regions, ticks are the most medically important group of vectors, transmitting a broad variety of pathogens. Warmer winters, longer humid seasons, and increasingly mild weather in northern latitudes have allowed ixodid ticks to push into latitudes and elevations they did not previously occupy, while urbanization concentrates interactions between competent reservoir hosts and human contact within the same gradients [1, 2]. In the USA, metropolitan New York City (NYC), lies near a humid subtropical climate (Cfa) and a humid continental climate (Dfa) boundary and sits at the leading edge of these changes [3]. It has extensive urban parkland, dense small and meso-mammal populations, a high introduction pressure [4], and multiple medically important tick vectors including the recently established Asian longhorned tick (*Haemaphysalis longicornis*). This tick species is cold-tolerant, parasitizes a wide range of animals, and reproduces parthenogenically allowing its rapid proliferation in the Northeast and Midwest United States after being first documented in 2017 [5, 6].

Ticks are vectors of many pathogens that cause debilitating diseases, and are transmitted during bloodmeal acquisition. Ticks frequently carry multiple pathogens simultaneously (co-infections) [7], complicating pathogen detection and disease treatment. In *Ixodes scapularis* (the blacklegged tick), one of the most prominent vectors in North America, these include bacteria *Borrelia burgdorferi*, *Borrelia miyamotoi*, *Anaplasma phagocytophilum*, *Ehrlichia chaffeensis*, Powassan virus (POWV), and the endoparasite *Babesia microti*. This vector has a multistage life cycle, feeding once before molting to the next life stage (or reproducing if adults). Different life stages may select hosts of different sizes. These hosts differ in their pathogen competence (i.e., ability of the host to successfully maintain and transmit pathogens to a vector) [8, 9]. In North America, white-tailed deer (*Odocoileus virginianus*) have been frequently implicated in the growth of tick populations because they are the most frequent host for adult blacklegged ticks, but aside from *Ehrlichia chaffeensis*, are generally considered non-competent for pathogen transmission [10, 11]. However, immature, blacklegged ticks can be found on all hosts, including deer [12]. Because of the difficulty in trapping hosts to determine their association with ticks, the ability to determine the previous bloodmeal of a host-seeking tick has been a fundamental goal of tick research. Linking tick infection status of host-seeking ticks to the bloodmeal host in the preceding life stage(s) indicates the relative roles of different hosts in the maintenance of pathogens and contributes to a broader understanding of vector-host interactions and complex pathogen transmission networks [4, 13].

Emerging viruses transmitted by ticks span 23 families, of which the *Flaviviridae* have broad host range and zoonotic potential [14]. This family is composed of two genomic classes: unsegmented and segmented. Tick-borne viruses (TBV’s) belonging to the *Flaviviridae* family are widely reported around the world and lead to more than 10,000 hospitalizations (Europe and Russia) annually [15]. Tick-borne encephalitis virus (TBEV) causes encephalitis, non-specific viremia, neurological, and hemorrhagic symptoms in the infected patients [15–17]. The primary known vectors for TBEV are *I. ricinus* and *I. persulcatus* [18, 19]. TBEV incidence is increasing in Europe and Eurasia [20–30], while Asian Omsk hemorrhagic fever virus (AOHFV), Louping ill virus (LIV), and Kyasanur Forest Disease (KFD) remain localized but globally distributed across Europe, Asia, and the Middle East. POWV is categorized as lineages I or II. POWV lineage II is steadily expanding in the Northeast and Midwest USA and is now considered an emerging pathogen with localized transmission foci [31, 32]. POWV illness is severe, with a fatality rate as high as 19% in adults [33]. Infection with segmented Jingmen tick virus (JMTV) has been detected in patients co-infected with fatal Crimean-Congo Hemorrhagic Fever virus (CCHFV) in Kosovo [34] and associated with mild to severe illness in patients infected in China [35]. The first report of a segmented TBV detection in the USA was in 2020 where rodents in Pennsylvania were identified with sequences with 70% amino acid identity with the JMTV polymerase, provisionally named *Flavi-like segmented virus US* (FLSV-US) [36].

Polymerase chain reaction (PCR) is the most used detection method in surveillance and is currently accepted as the standard. PCR is limited as it can only identify closely related pathogens, and when targeting highly conserved regions, it provides limited genomic information. This means that further analysis between closely related pathogens is required to determine taxonomical relationships [37]. Hybrid-capture Next Generation Sequencing (HCNGS) is an alternative enrichment method that addresses limitations of current metagenomic and amplicon-based sequencing methods. It requires biotinylated nucleic acid baits to complementarily bind to target DNA (or *in vitro* reverse-transcribed RNA) in prepared libraries during hybridization (Extended Data Fig. 1). Unlike PCR-based modalities, HCNGS allow for semi-targeted enrichment of targeted genomic regions in large quantities. A single HCNGS panel can hold hundreds to thousands of complementary baits and thus can facilitate the enrichment of many target sequences at once. Baits are generally 75 bp to 140 bp long and engineered with sequence degeneracy for variation tolerance. While the turnaround time and cost of HCNGS is still an area of development, it does not require any specialized equipment other than a benchtop Illumina sequencer, which is now increasingly common in public health and hospital laboratories [38–40].

Here, we report complementary hybrid-capture (HC) panels TITAN-RNA, targeting all known tick-borne viruses and key vector-borne febrile illness agents; TICKHUNTER, the companion bacterial, parasitic panel with an interchangeable bloodmeal panel. We demonstrate their integrated deployment for surveillance and potential for clinical diagnostics. We establish analytical performance against synthetic and naturally divergent targets; apply the panels to host-seeking ticks and meso-mammal samples collected across NYC and the Pacific Northwest; identify circulating *A. phagocytophilum* sequence types and a *Bab. microti*–like piroplasm in NYC raccoons; discover a chimeric segmented/unsegmented *Flavi-like virus* (FLV) in invasive NYC *H. longicornis*; and combine with codon adaptation analysis, serology, and bloodmeal mapping to define the urban transmission network of NYC.

## Results

### Analytical validation shows that the designed HC panels perform comparably to qPCR and maintain robust detection despite sequence variation

#### HC panel design

Two main panels were designed to detect and sequence TBVs (TITAN-RNA) and tick-borne bacteria and endoparasites (TICKHUNTER). For validation, different versions of each panel were designed, each containing different targets, and used on their own or in combination. TITAN-RNA v1 targets conserved genes from 19 common TBVs (Suppl. Table 2), including viruses known to circulate in the US (POWV) and those not reported in the US (CCHFV). TITAN-RNA v2 targets conserved genes of 91 novel and emerging TBVs reported worldwide (Suppl. Table 3). A final version, TITAN v3, combines all targets from TITAN-RNA v1 and v2, targeting the genes or genomes of 159 arboviruses (Suppl. Table 4). TICKHUNTER v1 targets 25 genes or genomic regions of common tick-borne pathogens (TBP), including bacteria like *Bo. burgdorferi* and parasites like *Bab. microti.* For this panel, highly conserved genes or genomic regions were included (Suppl. Table 5). To further characterize the detected pathogens, a second panel (TICKHUNTER v2) was designed to add 18 additional genes used for speciation and multi-locus sequence typing (MLST), and genes related to pathogenicity in certain species/genera (Suppl. Table 6). Finally, a panel for host blood meal identification was also designed (Blood Meal panel v1) and includes baits targeting a variable region of the mammalian mitochondrial 16S rRNA of nine mammalian species including humans (Suppl. Table 7).

#### The TITAN-RNA and TICKHUNTER panels show linearity and limit of detection comparable to qPCR

To assess the linearity of the HC panels, we used 10-fold serial dilutions of pooled synthetic dsDNA with the known sequences of eight and three pathogens for the TICKHUNTER V1 and TITAN-RNA V1, respectively. For the targeted pathogens, a linear relationship was identified between synthetic dsDNA copy number and number of identified reads resulting from HCNGS (Fig. 1A, and Extended Data Fig. 2), indicating suitability for quantitative analysis. To better approximate the complexity of clinical samples, the limit of detection (LOD) of the panels was tested with quantitative genetic material including genomic RNA of POWV and genomic DNA of *Bab. microti* spiked in water (n=6), blood (n=5), and ear skin biopsies (n=5) from laboratory mice (Fig. 1, Suppl. Table 9). The LOD using POWV genomic RNA in water (n=6) revealed that the HCNGS had a lower LOD (1.36 total copies) than the qPCR (9.82 total copies, Extended Data Fig. 2B). Probit regression modelling estimated an LOD for POWV genomic RNA of 3.24 total copies in ear skin biopsies and >5.625 total copies in blood (Fig. 1B). A complementary log-log regression model was used to assess skewed data and extrapolate an integer LOD of 36.6 total copies for POWV RNA in blood [41, 42]. For the genomic DNA of *Bab. microti*, the LOD was 87.4 total copies in ear skin biopsies, and 21.6 total copies in blood (Fig. 1B).

**Fig. 1.**
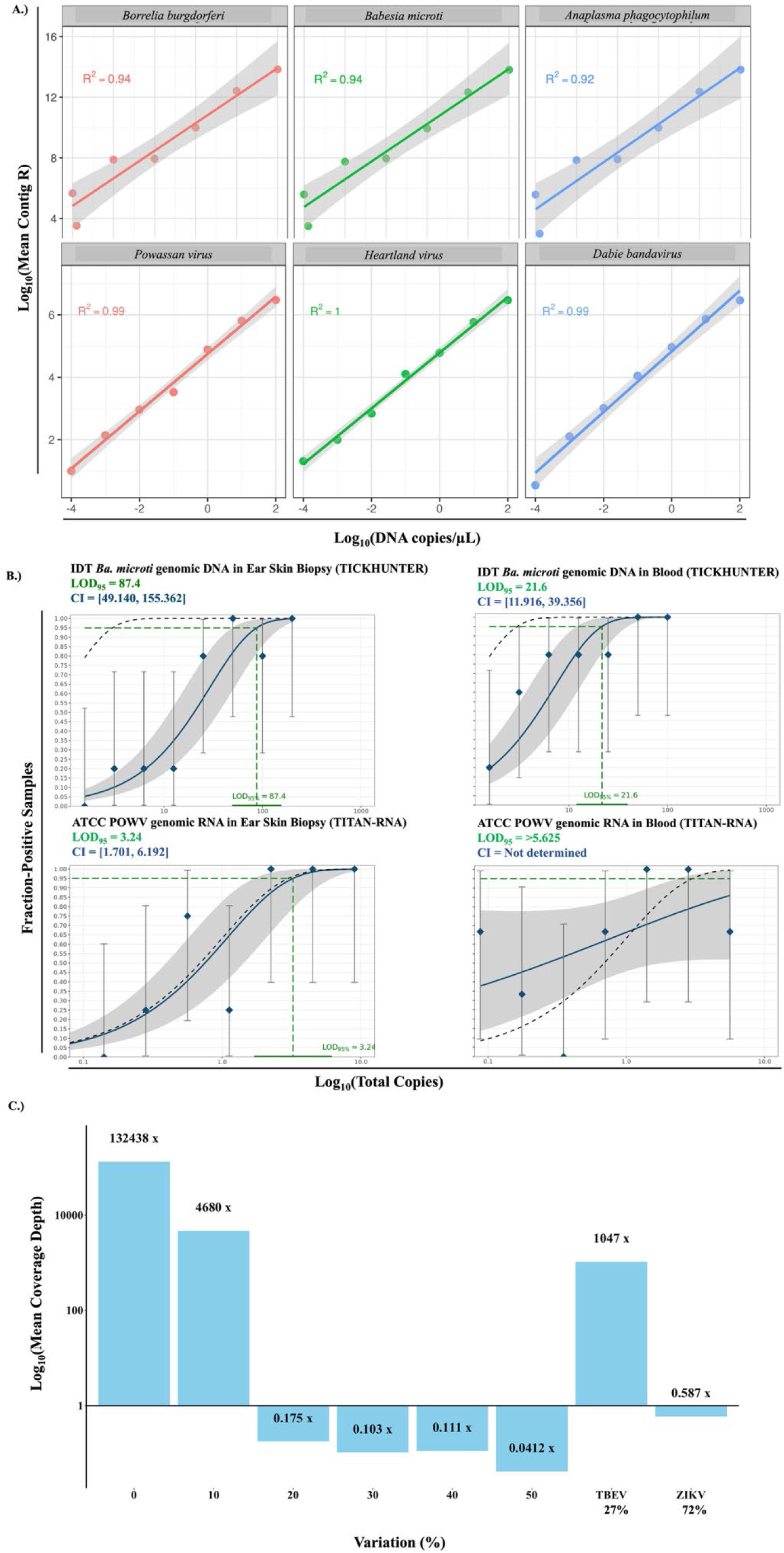
Analytic performance of the TITAN-RNA v1 and TICKHUNTER v1 HCNGS panels. A) Linearity graphs for three representative pathogens for each panel (TICKHUNTER v1 at the top, TITAN-RNA v1 at the bottom) plotted as Log10 of mean contig R (number of reads aligned per taxonomically matched contig, Y-axis) versus Log10 of synthetic dsDNA concentration (copies/µl, X-axis). The coefficient of determination (R^2^) is shown in the top left corner of each graph. The mean contig R was obtained after hybridization capture and sequencing of three and five replicates of each dilution series for the TITAN-RNA v1 and TICKHUNTER v1 panels, respectively. For the TICKHUNTER v1 panel, the linearity was tested for five additional pathogens (Extended Data Fig. 2). B) Fitted sigmoidal curves for the estimation of the limit of detection (LOD) of *Bab. microti* genomic DNA (top) and POWV genomic RNA (bottom) spiked in ear skin biopsy and blood samples from laboratory mice. C) Sequencing coverage plotted as Log10 of Mean Coverage Depth of POWV NS5 variants or wildtype POWV, TBEV, or ZIKV NS5 (n=9).

#### The TITAN-RNA panel tolerates genetic variation

To test the effect of natural and artificial variation in *Flaviviridae* in the detection and sequencing using the TITAN-RNA v1 panel, sixteen variants of the POWV NS5 gene were generated with 10, 20, 30, 40, and 50 percent random divergence. Two naturally divergent variants were also included (TBEV NS5 and ZIKV NS5 genes representing ∼27% and ∼72% natural variance, respectively) (Suppl. Table 8). The TITAN-RNA v1 panel, which includes baits targeting the NS5 gene of POWV lineages I and II, tolerated random sequence variation of up to 10%, yielding the highest coverage depth among the simulated variants. Natural sequence variability was tolerated up to 27% (TBEV), which produced the second-highest coverage depth, Fig. 1C). The baits did not reliably capture randomized synthetic sequences with variability higher than 10% and only ‘captured’ an average of < 10 ZIKV (72% variable) reads (Extended Data Fig. 3).

### The TICKHUNTER and Blood Meal panels detects and subtypes bacterial and parasitic targets and host bloodmeals

#### Combined HC panels can simultaneously detect multiple pathogens and host bloodmeals

For *Borrelia* detection in *Peromyscus leucopus* ear skin biopsies, real-time PCR and HCNGS agreed on 58 of 64 unique animals (four PCR-positive/HCNGS-negative, two PCR-negative/HCNGS-positive, Suppl. Table 11). Because observed prevalence differed between the two NYC study sites (Queens 17.9%, Staten Island 60.0%), we estimated performance without a reference standard using a two-population Bayesian latent class model: HCNGS Se median 91.7% (95% credible interval 72.2–99.6) and Sp median 97.7% (95% credible interval 89.6–99.9); real-time PCR Se median 93.1% (95% credible interval 76.3–99.5) and Sp median 94.1% (95% credible interval 83.3–99.5). From blood, although *B. burgdorferi* was recovered only from 2/39 mice tested by hybrid capture, *Bartonella ssrA* sequences were unexpectedly recovered from 25/39 blood samples, resolving two clades circulating in both sites: one with *Bar. rochalimae* (Group 1), and another closely related to *Bar. vinsonii, Bar. grahamii, Bar. japonica* and *Bar. coopersplainsensis* (Group 2). The sequences from Group 1 were identical and those from Group 2 were 99.8% similar. Each group contains sequences obtained from mice collected in both Queens and Staten Island (Fig. 2D).

**Figure 2.**
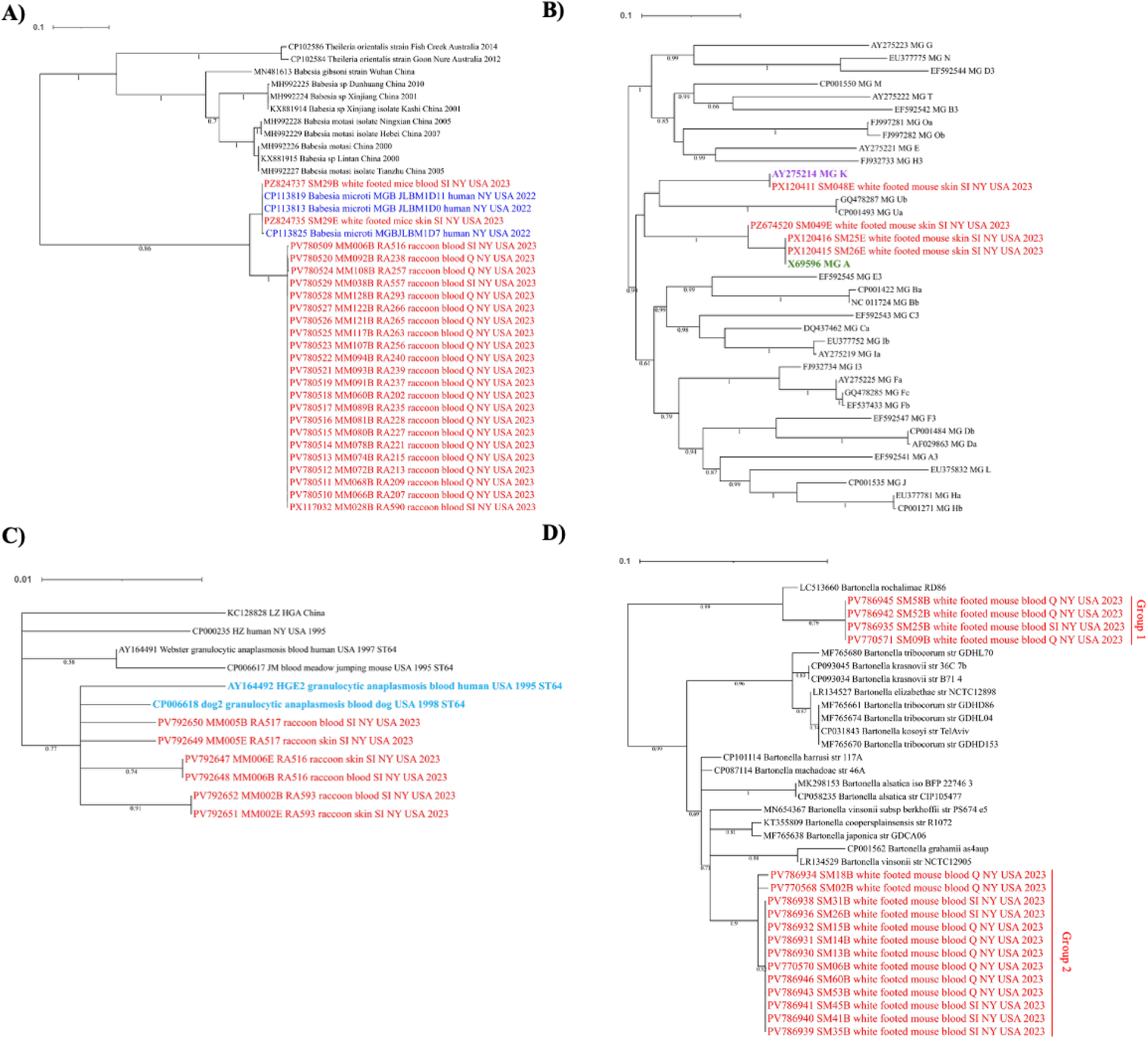
Phylogenetic analysis of tick-borne DNA pathogens detected in blood and skin biopsies of small mammals (SM) and meso-mammals (MM). Maximum likelihood (ML) phylogenetic trees of *Babesia* (A), *Borrelia* (B), *Anaplasma* (C), and *Bartonella* (D). All the sequences obtained in this study come from free-ranging white-footed mice (*Pe. leucopus*) or raccoons (*Pr. Loto*r) sampled in Queens (Q) or Staten Island (SI). For all the trees, the sequences obtained in this study are in red, numbers in the branches indicate bootstrap percentage values for 1,000 replicates, and branches with <50% support were collapsed. Each tree tip label includes its accession number, pathogen species (if known), host species (if known), and collection site and year. NCBI accessions from this study in Suppl. Table 16. **A)** ML phylogenetic tree of a partial region (2,206 nt) of the 23S rRNA gene of selected *Babesia* species. The 22 sequences obtained in this study from raccoons form a group closely related but separate from the sequences from three *Bab. microti* reported in humans in New York State in 2022 (in blue). The sequences from two *Bab. microti* obtained from a white-footed mouse group within the *Bab. microti* group. Nucleotide substitution model used: GTR+G+I. **B)** ML phylogenetic tree of a partial region (481 nt) of the *ospC* gene of 33 major groups (MG) of *Bo. burgdorferi* sensu stricto. Three sequences obtained in this study from white-footed mice from SI group within MG A (in green), and the fourth one groups with MG K (in violet). Nucleotide substitution model used: GTR+G+I. **C)** ML phylogenetic of the partial sequence of the msp2-B gene of *A. phagocytophilum* (465 nt). The six sequences obtained in this study group together with members of the allelic group ST64 detected in the blood of a human and a dog with granulocytic anaplasmosis (in light blue). Nucleotide substitution model used: HKY+G. **D)** Maximum likelihood phylogenetic tree of a 242 nt region of the *ssrA* gene of *Bartonella*. The seventeen sequences obtained in this study fall into two groups. Both groups include sequences detected in blood samples from white-footed mice in Q and SI, NYC. The species of *Bartonella*, strain (str), and isolate (iso) are shown in the sequence name, when known. Nucleotide substitution model used: K2+G.

Eight pools of *I. scapularis* larvae removed from eight individual mice were tested as proof of concept to assess the simultaneous detection of host bloodmeal and tick-borne pathogens using the TICKHUNTER v1 and the Blood Meal panel v1 combined. *Pe. leucopus* mitochondrial DNA was detected in all pools. Simultaneous detection of on to three pathogens from the larvae in addition to the *Rickettsia* endosymbiont of *I. scapularis* were also demonstrated. Pathogenic gene sequence coverage was as high as ∼6,600X from a single larva and ∼133,500X from 2 pooled larvae (Suppl. Table 15).

#### The TICKHUNTER panel enables pathogen typing from blood and skin biopsy samples

Subtyping directly from animal (mouse or raccoon) specimens with low-pass (iSeq) sequencing (511,418 to 846,326 total reads per sample) was performed for *Bo. burgdorferi*, and medium-pass (MiSeq, 190,612 to 2,052,102 total reads per sample) was performed for *Babesia*, *Anaplasma*, and *Bartonella* species to simulate point-of care testing conditions. A segment of the 23S rRNA of *Babesia* spp. was detected and sequenced from blood in 22/24 raccoons. The *sa1* gene of *Bab. microti* was not detected in any sample, either by qPCR or HCNGS (Suppl. Table 11). The obtained 23S rRNA sequences shared 99.98% similarity with each other and clustered together in a single group (Fig. 2A). This group was closely related to three *Bab. microti* strains (1D0, 1D7, and 1D11), identified from human cases in New York State, USA, in 2022. Genetic comparison of these three *Bab. microti* with the 22 raccoon sequences obtained in this study showed that they are 98.2% similar and appear to be an unexpected *Bab. microti*-like species in which *sa1* was not detected. In contrast, the *Bab. microti* detected in white-footed mice grouped within the *Bab. microti* group (Fig. 2A), and the *sa1* gene was detected in all the positive white-footed mouse samples (Suppl. Table 11).

The TICKHUNTER v2 panel targets 16 genes in total for the MLST of *A. phagocytophilum* and *Bo. burgdorferi.* From the seven genes targeted for the MLST of *A. phagocytophilum* [43], we obtained complete or partial sequences from blood and ear skin biopsies of three raccoons (RA516, RA517, RA593). Fewer genes were detected and sequenced from ear skin biopsies, with only one to three genes sequenced, compared to blood samples where a minimum of six genes were recovered (Suppl. Table 12). Based on the more comprehensive results from blood, the closest allelic group (or ST) for the samples detected from two raccoons (RA516, RA517) was ST64, and for the third raccoon (RA593) was ST161. In a phylogenetic tree using the longest concatenated MLST sequences obtained for two of these samples (RA516 and RA593), these strains clustered with three other allelic groups (ST64, ST161, ST215), all of which have been reported in the blood of humans or horses with granulocytic anaplasmosis in the USA between 1969 and 2007 (Extended Data Fig. 4). We obtained additional partial or complete sequences of the *msp4, msp2-B,* and *msp-2C* of *Anaplasma* from the same three raccoons (Suppl. Table 13). For two of the sequences (one of *msp-2C* and one of *msp2-B*), we sequenced the complete gene, plus flanking sequences. As with the MLST genes, longer and more complete sequences were detected in whole blood than in ear skin biopsies samples. However, it was possible to obtain sequences, complete or partial, for all the samples, facilitating sequence comparisons between sample types. The partial sequences of *msp2-B* (465 nt) from the same individual were identical, while a comparison between individuals showed that the sequences were 99.8% similar. A phylogenetic tree of this partial region of the *msp2-B* gene for both whole blood and ear skin samples for the three raccoons revealed their grouping with those from the allelic group ST64 (Fig. 2C), as seen using the MLST genes (Suppl. Table 12).

From the nine genes targeted for the MLST scheme of *Bo. burgdorferi*, a minimum of four and a maximum of seven genes were recovered in 18/22 positive samples (Table 1), including 10 samples with putatively low spirochete loads (real-time PCR Ct range 28.6-35 from 10 samples), at 511,418 to 2,805,550 reads per sample. Although genome coverage declined as Ct values increased, complete (100%) coverage of certain genes remained achievable. This was accomplished with an average of 629,420 ± 90,881 reads per sample in a low-pass sequencing platform (iSeq) with Ct values of 28.6-32.4, and 2,286,126 ± 299,754 reads in a medium-pass platform (MiSeq) with Ct values of 28.6 to 34.5 (Table 1). We obtained the complete sequence of *ospC* for 4/10 skin biopsy samples that tested positive for *Bo. burgdorferi* with high Ct values. A phylogenetic tree constructed from these sequences, along with those from 33 major groups (MG) of *Bo. burgdorferi* sensu stricto, indicates that three sequences belong to MG A and one to MG K, with potential for invasive human disease (Fig. 2B) [44, 45].

**Table 1.** Summary of *Borrelia burgdorferi* read counts required for low and medium-pass Illumina sequencing for direct pathogen subtyping from skin biopsies.

| Mouse ID | Real-time PCR Ct | Low-pass sequencing platform (iSeq) |  |  |  | Medium-pass sequencing platform (MiSeq) |  |  |  |
| --- | --- | --- | --- | --- | --- | --- | --- | --- | --- |
|  |  | Fraction of targeted genes sequenced | % min coverage | % max coverage | Total number of reads | Fraction of targeted genes sequenced | % min coverage | % max coverage | Total number of reads |
| SM48E | 28.6 | 100 | 85.8 | 100 | 573,976 | 100 | 100 | 100 | 2,001,200 |
| SM45E | 30.3 | 100 | 41.4 | 100 | 756,858 | 100 | 61 | 100 | 2,616,534 |
| SM25E | 31.1 | 100 | 80 | 100 | 555,528 | 100 | 94.65 | 100 | 2,141,678 |
| SM26E | 32.1 | 100 | 30.7 | 93.4 | 788,486 | 100 | 89.8 | 100 | 2,739,836 |
| SM64E | 32.4 | 100 | 31.8 | 100 | 631,316 | 100 | 64.6 | 100 | 2,328,880 |
| SM65E | 32.4 | 14 | 26.4 | 26.4 | 556,970 | 28 | 26.4 | 63.8 | 1,715,320 |
| SM35E | 33.2 | 71 | 28.6 | 90.3 | 846,326 | 100 | 43.7 | 94.6 | 2,805,550 |
| SM42E | 33.6 | 100 | 29.4 | 95.8 | 619,066 | 100 | 38.5 | 100 | 2,230,930 |
| SM49E | 34.5 | 71 | 27.6 | 99.2 | 511,418 | 100 | 45.8 | 100 | 1,943,822 |
| SM43E | 35.0 | 57 | 35.3 | 62.7 | 698,140 | 100 | 22.6 | 92.1 | 2,465,146 |

#### A segmented/unsegmented Flavi-like virus in invasive *H. longicornis*

A total of 216 pools of host-seeking *H. longicornis* pools collected by community volunteers in the NYC area were screened with TITAN-RNA v1, and 23 pools screened with v1 and v2 combined (Fig. 3A). Ticks were pooled by species, life stage, and location collected. Homogenates with any detection of exotic *Flaviviridae* were referred for deep untargeted Illumina RNAseq at an academic core facility and supplemented with additional targeted long-read amplicon sequencing. Three pools revealed VP1ab-like glycoprotein sequences of non-endemic Flavi-like Segmented Viruses (FLSV’s, Fig. 3B and 3D). Alignment and phylogenetic analysis revealed that sequences from pooled samples FLSV-NY-2022-36 (330 bp), FLSV-NY-2022-44 (402 bp), and FLSV-NY-2022-93 (527 bp) clustered closest to *Jingmen tick-like* viruses. FLSV-NY-2022-36 contained five larvae, FLSV-NY-2022-44 five nymphs, and FLSV-NY-2022-93 five nymphs. The TITAN-RNA Panel v1 did not target any FLSV glycoprotein sequences, and we hypothesize that our initial detection of unsegmented *Flavivirus* glycoprotein sequences was likely due to the high flexibility probes binding to FLSV glycoprotein sequences later revealed by bulk RNA sequencing. TITAN-RNA Panel v3 was modified to target the FLSV-NY glycoprotein sequences a dectected a longer putative glycoprotein sequence recovered five fresh bovine host-removed *H. longicornis* obtained from the Northern edge of the infestation in May 2026 (Columbia County, NY, location not shown to protect privacy). This sequence is identical to *Alongshan virus* (ALSV, NCBI# MZ676705) and *Cheeloo Jingmen-like virus* (CJLV, NCBI# OR114880.1), and the partial glycoprotein sequences recovered from the host-seeking NYC ticks match the sequences recovered from the bovine-removed.

**Figure 3.**
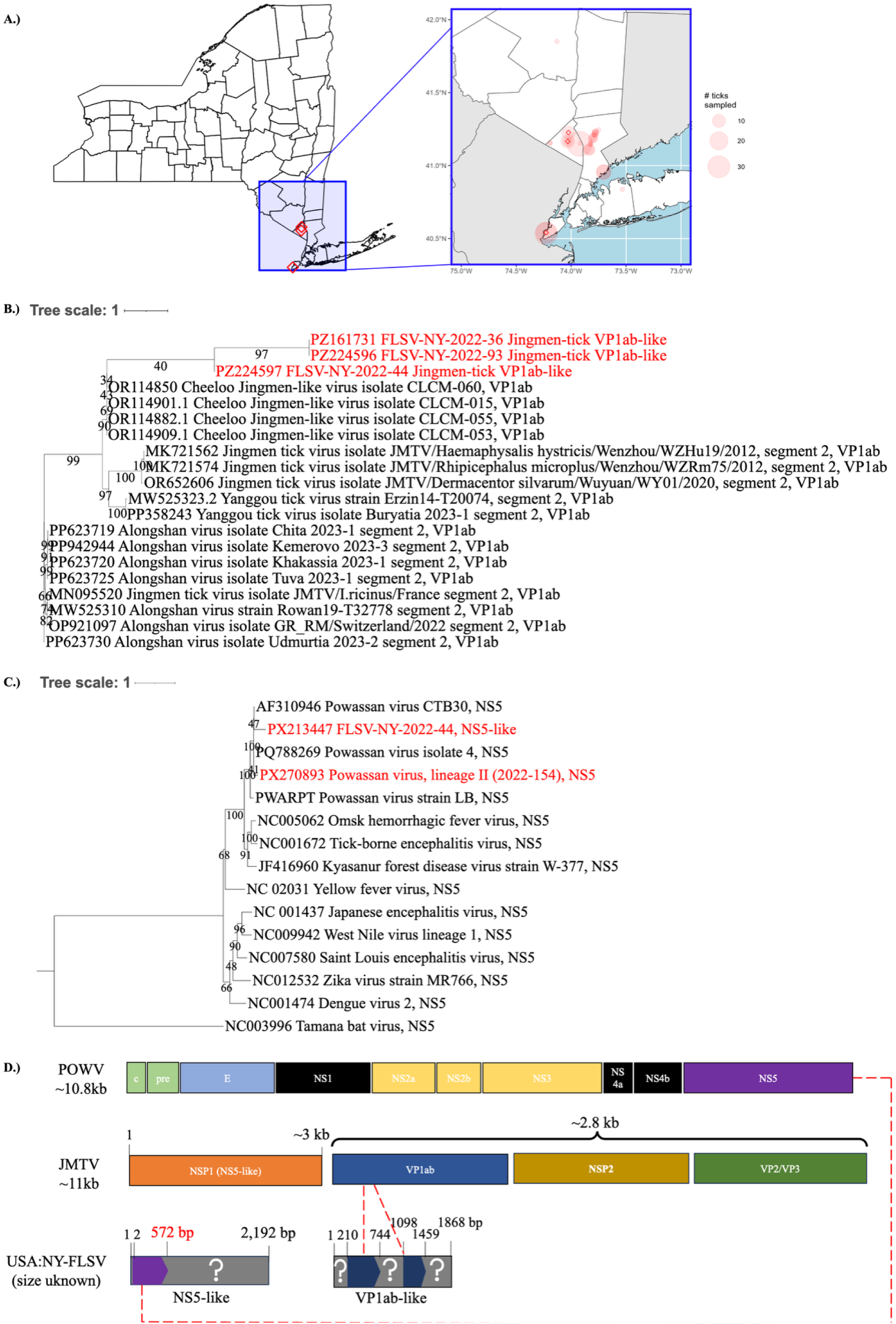
Evolutionary support that FLSV-NY is a chimeric Flavi-like virus. A) Geospatial mapping of ticks sampled in NYS. Collected ticks are plotted over their sampling centroid with jitter and scaled to represent total number of samples collected. B) Phylogenetic analysis of tick-borne *Flavivirus* NS5. Maximum likelihood tree of unsegmented *Flavivirus* NS5 homologs. *Tamana bat virus* was used as the outgroup [49]. Scale represents number of nucleotide substitutions. Bootstraps = 1000. NCBI accessions from this study in Suppl. Table 16. C) Phylogenetic analysis of tick-borne *Jingmenvirus* glycoprotein (VP1ab). Maximum likelihood tree of segmented *Jingmenvirus* glycoprotein homologs. Scale represents number of nucleotide substitutions. Bootstraps = 1000. NCBI accessions from this study in Suppl. Table 16.

To assess the broader distribution of *Jingmen*-like glycoprotein sequences beyond the three FLSV-NY-positive discovery pools, we developed a qPCR (JMLV) assay targeting the FLSV-NY/ALSV glycoprotein gene. The bias-corrected maximum-likelihood prevalence (EPP) in New York *H. longicornis* was 97.3% (95% CI 84.7–100%, n=265 pools/1,171 ticks) for host-seeking and 94.1% (CI 84.4–98.7%, n=58 pools/114 ticks) for host-attached specimens (Suppl. Table 19). JMLV was largely restricted to this species; host-seeking *A. americanum* (0/3) and host-attached *I. scapularis* (0/13) and *I. cookei* (0/1) were entirely negative, while host-attached *A. americanum* (1/79) and host-seeking *I. cookei* (1/7) each returned a single isolated positive against an otherwise negative background.

One of the samples (FLSV-NY-2022-44) had both a 402 bp contig matching the glycoprotein gene of segmented *Jingmen tick virus* and an approximately 2.2 kb highly modified sequence putatively from an unsegmented *Flavivirus* NS5 (Figs. 3C and 3D). The putatively recombinant NS5 sequence branched closest to POWV CTB30 (NCBI# AF310946.1). A 570 bp region of the sequence had 90.49% nucleotide identity (29% query coverage) to the NS5 of the clinical POWV isolate, while the remaining sequence was highly divergent, with 30.3% amino acid identity (9% query coverage) to an exogenous uncharacterized protein in *H. longicornis* (NCBI# KAH9380833.1). Complementary regions of approximately 26 to 33 bp flank the 5’ and 3’ ends of the recovered sequence, suggesting the ability to form a hairpin-like structure putatively necessary for viral replication.

Mapping of RNAseq to mitochondrial 16s rRNA (*mt16s*) and confirmation with cytochrome *B* (*cytB*) [46] revealed a low abundance of *P. lotor* (NC_009126.1)*, Canis* spp., and *Bos* spp. Specific sequences, suggesting them as putative blood acquisition hosts (Extended Data Fig. 5A and 5B). Hypothesizing that FLSV-NY would be cross-reactive in a crude antigen TBEV ELISA (as it was likely captured by those HC probes), we screened the banked 2023 sera from 188 NYC mammals (128 raccoons, 11 skunks, 5 opossums, and 44 mice) with a commercial ELISA kit alongside PCR results from 153 host-attached ixodid ticks collected in 2023-24. Skunks had concentrations ranging from 0 to 281.9 U/mL, opossums from 0 to 77.8 U/mL, and mice from 0 to 88.1 U/mL (Fig. 4B). Raccoon sera had significantly higher antibodies binding to TBEV antigen than mice, with antibody concentrations ranging from 4.3 U/mL to 472.3 U/mL (median 59.8 U/mL), suggesting variation in immune regulation or population exposure. Raccoon serology was statistically analyzed with Kruskal-Wallis with Dunn’s post-hoc (Padj < 0.001, Fig. 4B). Staten Island and Queens had the highest populations of TBEV reactive raccoons compared to other boroughs (Fig. 4C).

**Figure 4.**
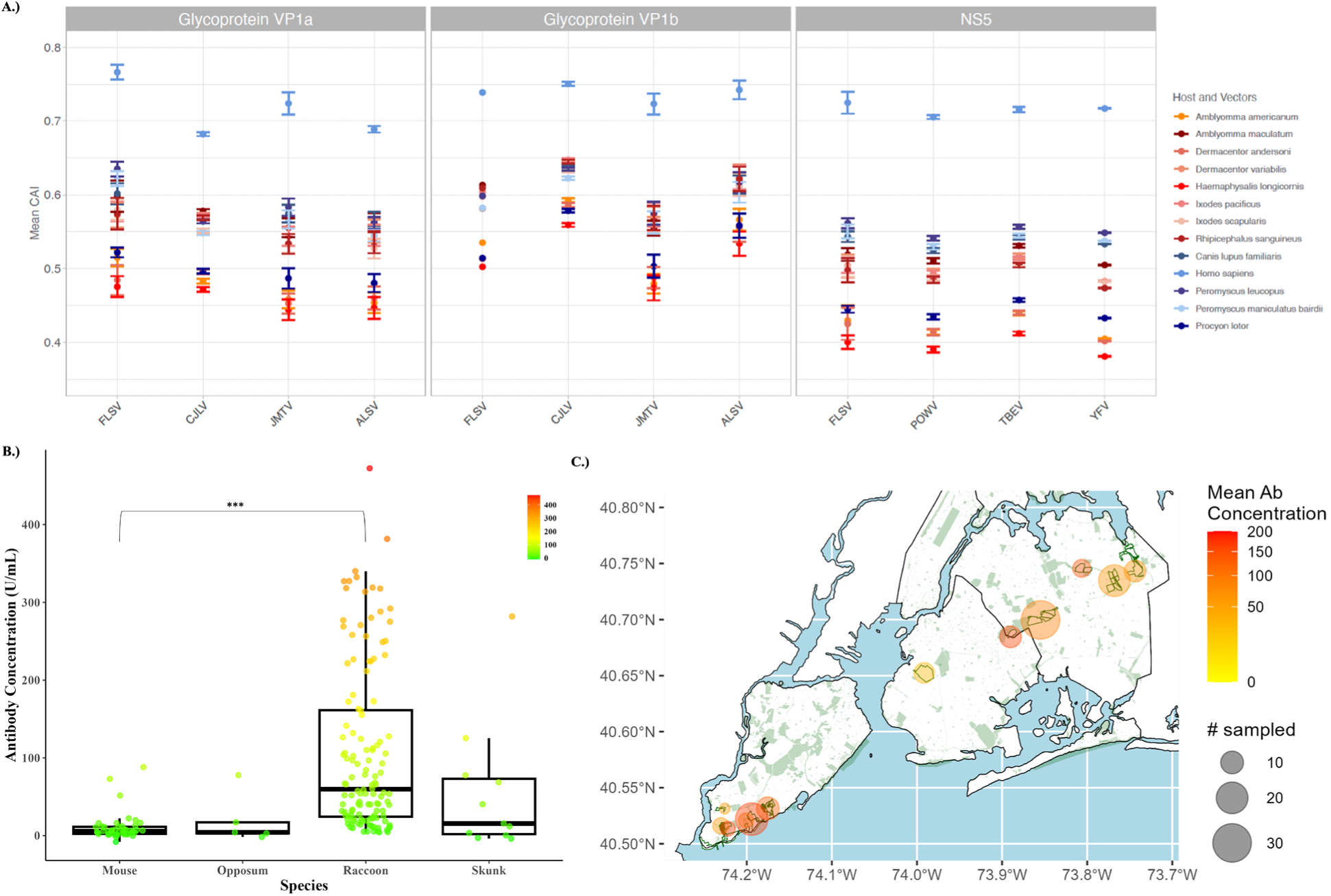
Raccoons are a putative reservoir host for FLSV-NY. A) Codon adaptability index (CAI) analysis for FLSV-NY and closely related viruses based on the codon usage of the most frequent genes of host and vector species. CAI values based on glycoprotein *vp1a* and *vp1b* of FLSV-NY as well as segmented viruses CJLV, JMTV and ALSV; and NS5 of FLSV-NY as well as unsegmented viruses POWV, TBEV and YFV reflect potential adaptation of these viruses to human, dog and prairie deer mouse (*Peromyscus maniculatus bairdii*) as hosts, and several North American tick as vectors. Values are reported as mean ± standard deviation. B.) Antibody concentration by species (TBEV ELISA). Antibody concentration in U/mL detected in mouse, opossum, raccoon, or skunks in NYC. The sample sizes for skunks and opossums were too small to determine significance. C.) Antibody concentration by borough. Antibody concentration in U/mL detected raccoons in Brooklyn (BK), Queens (Q), or Staten Island (SI).

#### Codon adaptation of the FLSV-NY indicates human compatibility

To provide a preliminary indication of the virus’s potential to infect humans, a range of secondary vectors, and hosts, we investigated FLSV-NY’s adaptation to North American tick vectors, human, and other mammalian hosts by comparing the codon adaptation index (CAI) estimates based on the codon usage bias of its glycoprotein VP1a and VP1b, and NS5 genes (Fig. 4A; [47]). Codon adaptation to different host and vector species varied between segmented viruses (FLSV-NY, CJLV, JMTV, ALSV) and non-segmented *flaviviruses* (POWV, TBEV, YFV). For all three viral genes analyzed (Glycoproteins VP1a, VP1b, and NS5), the tick-borne segmented *flaviviruses* (especially ALSV and JMTV) displayed moderate CAI values (0.5–0.6 range) across most tick vectors, including *A. americanum*, *I. scapularis*, and *Rhipecephalus sanguineus*, suggesting broad adaptation to multiple arthropod species. In comparison, CAI values for FLSV-NY were generally lower across vertebrate animal or invertebrate hosts in the NS5 gene.

All genes displayed high CAI values for *H. sapiens* and *C. lupus familiaris* as compared to reservoir/intermediate hosts or vectors, indicating possible codon optimization in these hosts. Across the board, we see similar trends among vertebrate hosts with NS5, glycoprotein VP1a and VP1b showing high CAI values for *Pe. leucopus* and *P. maniculatus* are higher, closely followed by *C. lupus familiaris* and with *P. lotor* showing the lowest CAI values. In ticks, NS5, glycoprotein VP1a and VP1b express the highest CAI values for *A. maculatum*, closely followed by *R. sanguineus*. *I. pacificus*, *I. scapularis*, *Dermacentor variabilis*. *H. longicornis*, *D. andersoni*, and *A. americanum* have the lowest CAI for these proteins. In general, CAI estimates were higher for mammalian hosts than ticks, with exemption of *P. lotor*, whose CAI values were more like ticks. All CAI findings are consistent with analysis performed using the Puigbò et al. eCAI web application [48], and results can be found in the source data for Fig. 4A.

#### Bloodmeals reveal potential *Powassan virus* and Tick-borne Relapsing Fever spirochete hosts

Bloodmeal analysis with mitochondrial 16S rRNA metabarcoding on six POWV positive, host-seeking *I. scapularis* samples (nymphs or adults) revealed that in addition to deer, dogs, cats, and birds may have been fed upon in this landscape; host competence was not assessed (Extended Data Fig. 5C). *Odocoileus virginianus* was the sole bloodmeal remnant detected in one host-seeking nymph pool with POWV Ct 27 (2022-154) collected in Nassau County, NY by community volunteers [50]. Typed as lineage II with TITAN-RNA, the closest relative based on the NS5 sequence (43 reads) was a strain from Centre County, PA (NCBI # 8269) (Fig. 3C). Pools with qPCR Ct range 36.1-42.3 could not be subtyped. In two of six lineage II-infected adults collected in the same county in 2018 [31, 51], *Canis lupus familiaris* (domestic dog) was the presumed nymphal bloodmeal remnant based on predominant relative abundance. A mammal from *Felidae* spp. was the predominant remnant in two adults and a possible larval bloodmeal source for one of these. *Haemorhous mexicanus* (house finch) was the presumed larval meal in one of these adults (LI108).

As a proof of principle for identifying bloodmeal sources to guide serosurveillance in areas with unidentified human illnesses, we investigated a collection of *Ornithodoros hermsi* (four adults and three nymphs), from premises outside of our primary study’s temporate zone in the Pacific Northwest USA with cases of unexplained relapsing fever, both with HCNGS-targeted host detection and host-agnostic mitochondrial 16S rRNA metabarcoding. Bloodmeal analysis of these soft ticks detected feeding on mice, chipmunks, bats, domestic dogs, cats, and horses (Extended Data Fig. 5D). For hosts that were included in the pilot panel, there was genus-level agreement between metabarcoding and HCNGS. No single method yielded more consistent results, although the host-agnostic approach provided additional resolution. No pathogens were detected using the pilot pathogen panels. Metadata for the aforementioned ticks can be found in Supplemental Table 14.

## Discussion

The possible weather-driven range expansion of ticks and their co-infection with pathogens is opening a diagnostic gap that single antigen-based assays were not designed to close. Hybrid capture, with its tolerance for sequence divergence and capacity to multiplex thousands of targets in a single workflow, is structurally suited to this problem. We show here that HCNGS with two complementary panels, TITAN-RNA and TICKHUNTER, matches qPCR for sensitivity and dynamic range, exceeds it for variant tolerance (up to 27% natural divergence), and together returns both a diagnostic answer and the genomic data required for evolutionary monitoring.

In this study, the TITAN-RNA v1-3 and TICKHUNTER v1-2 were run separately, and reverse transcription (RT) was only performed for the virus experiments. The slightly better LOD for RNA targets with HCNGS compared to qPCR may have resulted from performing the RT independently from PCR amplification. Performing a standalone RT for the qPCR may have improved its performance. Moreover, performing RT on samples with DNA targets may improve performance for the bacterial and parasite targets by capturing microbial cell-free DNA as well as replicating (mRNA or rRNA) pathogen genomes as demonstrated for *Bo. burgdorferi* by Branda et al. and reviewed more generally by Hu et al. [52, 53]. Future directions include combining the panels, adding additional diagnostic targets as they are discovered, and subjecting all samples to RT. All panels reported here are available for custom ordering through Twist Biosciences.

The highest coverage of variant *Flaviviridae* sequences by HCNGS was, as expected, observed in the regions with high similarity and nucleotide conservation. Coverage of less conserved sequences was attributed to both the flexible complementary binding action of the baits during hybridization and the pulldown of flanking sequences that does not depend on similarity to bait sequences. Variation tolerance was further demonstrated using field samples of invasive ticks found to harbor a potentially novel virus species combining segmented and unsegmented flavivirus genomes, with no *a prori* knowledge of its existence. The NS5 sequence was only recoverable with hybrid assembly and an extended reverse transcription, supporting its identity as part of a viral RNA genome rather than derived from the tick or a bloodmeal host genome. It is not related to the “FLSV-US” NS5 sequence discovered in mice in Pennsylvania, USA [36].

The detected FLSV-NY sequences exhibited codon adaptation across its glycoprotein VP1a and VP1b, particularly in *H. sapiens*, where CAI values exceeded those of other segmented TBVs. This elevated adaptation suggests a potential compatibility with human codon usage, raising the possibility that humans could act as hosts, though further investigation is required to assess replicative capacity in human tissues. In addition to humans, overlapping CAI values for FLSV-NY were observed in *C. lupus familiaris* and *P. maniculatus bairdii*, indicating similar levels of codon optimization across these mammalian hosts. Among arthropod vectors, FLSV-NY VP1a and VP1b genes showed moderate CAI values in *R. sanguineus*, *D. variabilis*, and *I. scapularis*, with *D. andersoni* consistently displaying the lowest CAI estimates across all three viral genes. This CAI pattern was broadly shared by other viruses in the panel (CJLV, JMTV, ALSV), suggesting conserved codon adaptation trends among tick-borne flaviviruses. For the non-structural gene NS5, FLSV-NY showed moderate to high CAI values across hosts, again with *H*.

*sapiens* ranking highest. Overlapping host-vector groupings (*C. lupus familiaris*, *P. maniculatus bairdii*, *R. sanguineus*, *D. variabilis*, *I. scapularis*) were observed, reinforcing a recurrent codon adaptation structure across viral genes. Overall, the FLSV-NY sequences appear to demonstrate greater codon optimization in mammals than in ticks, with its structural genes showing stronger host-specific adaptation, potentially indicating enhanced vertebrate compatibility. The adaptation results are based on short sequences and should be interpreted with caution. Brown et al. (2024) recently highlighted the presence of uncharacterized protein sequences that may code for endogenous or exogenous RdRP’s in the NCBI database and the need to annotate sequences towards higher accuracy for virus discovery [54]. The sequences identified using the TITAN-RNA panel will be added to future builds to detect additional sequence for more comprehensive evolutionary analyses. Available whole genomes of segmented and unsegmented members of the *Flaviviridae* family will be added, with a focus on glycoproteins (E and VP1ab) and RdRP (NS5) homologs. The TITAN-RNA v1 and TITAN-RNA v2 panels can be ordered through Twist Biosciences as one combined panel (TITAN-RNA v3), now including the novel sequences reported here and viruses from other arthropods. Combining the panels minimizes cost and labor towards an accessible method that can be adopted in hospital and academic laboratories around the world.

Ticks are responsible for 77%-95% of vector-borne disease cases in the US, and globally, TBVs are more prevalent in other parts of the world outside the US [55–57]. From an ecological perspective, the prevalence of TBVs may increase with land use changes, environmental change and increased host populations [29, 30]. Since tick-borne pathogens primarily infect small mammals, pathogen surveillance is necessary on small synanthropic mammals as they may serve as pathogen reservoirs and increase the risk of zoonotic spillover. As proof of principle, sera obtained from meso-and small-mammals in parks in Staten Island, Queens, or Brooklyn, NYC revealed antibodies capable of binding TBEV were detected in raccoons, skunks, opossums, and mice via crude antigen ELISA. Staten Island and Queens raccoons had the highest seropositivity. TBEV has not been reported in the US and its virion structure, which is highly conserved within *Flaviviridae* is likely targeted by antibodies specific to other *Flaviviruses* [58–68]. As aforementioned, three invasive *H. longicornis* ticks in this study were found containing putatively novel *Jingmen-like virus* sequences one being collected in Staten Island, and the others in Rockland County, NY. Our combined serological and detection results might suggest that well-known or putatively novel viruses belonging to *Flaviviridae* are circulating in NYC, particularly in Staten Island and Queens [67]. The presentation of seropositivity in raccoon sera suggests that they may serve as a reservoir host for tick-borne or other arthropod-borne *Flaviviruses*. If raccoons sustain bloodstream replication of *Flaviviridae* (and zoonotic strains of *A. phagocytophilum*) at population scale, the’dilution host’ framing of urban mesocarnivores may be incorrect, or more nuanced than previously thought. Future work will include screening suspected viremic raccoon sera for *Flaviviruses* with target-enrichment sequencing. As expected of *H. longicornis’* wide host range [69, 70], bloodmeal analysis revealed canines, raccoons, and potentially cattle as putative blood acquisition hosts for the segmented Flavi-positive ticks. Human DNA was also detected but given the nature of sampling cannot be distinguished as bloodmeal remnant or environmental contamination.

Unsegmented flaviviruses like TBEV, LIV, AOHFV, KFD, and POWV are harbored by ticks and well characterized genomically. Genomic references for more recently discovered segmented *Flavi-*like *jingmenviruses* are in development [71–74]. Our results highlight the need for increased monitoring of emerging disease vectors, including *I. scapularis* and the invasive *H. longicornis* species in the eastern USA, both competent vectors of POWV [31, 75–77]. Segmented and unsegmented viruses belonging to the *Flaviviridae* family can cause severe disease such as encephalitis, meningitis, loss of coordination, speech difficulties, high fever, and headaches [35, 78, 79]. Continued development of flexible enrichment methods for high-throughput sequencing technology will allow for improved detection and characterization of endemic and emerging viruses, potentially ahead of their spillover to humans.

*Bab. microti* is the most significant tick-borne intraerythrocytic protozoan causing human babesiosis in the US [80]. A higher diversity of *Babesia* spp. and related piroplasms has been reported in wildlife and certain human populations in specific regions [81–85]. The *Bab. Microti* surface antigen 1 (sa1) is a secreted protein critical for parasite invasion of host red blood cells, making the *sa1* gene a typical key target for detection [86]. The *sa1*-negative species identified here in blood from 22/24 raccoons showed remarkable similarity (99.98%) and clustered with three *Bab. microti* sequences from human cases in New York State in 2022 (Fig. 2A). *Bab. microti*-like species were reported by Garrett et al. [87] in raccoons from the southeastern US. The results further highlight the ability of the HCNGS method to identify unexpected but potentially significant taxa that are not identical to the baits, which can then be investigated further with targeted screening to assess their pathogenic potential.

MLST is a valuable tool for understanding the genetic diversity and population structure of pathogenic microorganisms by analyzing sequence variations in multiple housekeeping genes, making it an important tool for epidemiological surveillance [88–92]. Our approach allows for the detection of diverse gene sequences in a single workflow without the need for complex primer development needed for multiplex PCR [93]. *A. phagocytophilum* is a Gram-negative, obligate intracellular bacterium that primarily infects neutrophils, leading to granulocytic anaplasmosis in humans and various animals [87]. It is transmitted by *Ixodes* tick species, and has a broad host range, including humans, domestic animals, and wildlife. In the US, *I. scapularis* and *I. pacificus* are responsible for its transmission [91]. The bacterium’s ability to infect multiple hosts and its widespread distribution make it a significant concern for both veterinary and public health [87]. In this study, we targeted seven housekeeping genes of *A. phagocytophilum,* which encode proteins essential for maintaining fundamental cellular functions to ensure accuracy in detecting this bacterium [43, 93]. Additionally, we included more specific targets e.g. *msp2-B/msp-2C* and *msp4*, which encode polymorphic major outer surface proteins involved in erythrocyte invasion, to enhance the sensitivity of our approach [92]. These findings not only highlight the strengths of the HCNGS method in identifying genetic diversity in *A. phagocytophilum* but also provide valuable insights into its phylogenetic relationships and zoonotic potential. By revealing that the detected *A. phagocytophilum* is distinct from the Ap-V1 strain, which is thought to be non-pathogenic for humans [94], but is associated with ST64 (Fig. 2C), the use of our method will contribute to a better understanding of the epidemiology of granulocytic anaplasmosis.

Comparable detection of MLST genes was achieved across the two smallest Illumina benchtop platforms currently available (the iSeq and the MiSeq), and positive identifications required only tens to hundreds of reads per target. This read economy is a durable practical advantage. As platforms evolve and current legacy benchtop systems are succeeded by newer instruments, the workflow remains viable on whatever entry-level platform is current. Hybrid capture’s combination of high multiplex per run and low read demand per identification translates directly into lower per-sample cost and faster turnaround than metagenomics. The HCNGS method could therefore be deployed for rapid resilient assessment of tick-borne disease risk in economically underserved regions, regions of military deployment, and rural clinics handling unfamiliar febrile illness.

Two practical implications follow for biosurveillance in vectors as weather systems evolve. First, our results highlight the need for increased monitoring of emerging pathogen vectors, including *I. scapularis* and the invasive *H. longicornis* species in the eastern USA. Second, continued development of flexible enrichment methods for high-throughput sequencing technology will allow for improved detection and characterization of endemic and emerging pathogens, ideally ahead of their spillover to humans. Ultimately, surveillance and diagnostics paradigms should shift from a retrospective accounting of disease into a forward-looking and environment-agnostic early warning system.

## Methods and Protocols

### HC panel Curation and Design

#### Target curation

Two main panels were designed for the detection and sequencing of tick-borne viruses (TITAN-RNA) and tick-borne bacteria and endoparasites (TICKHUNTER). For the TITAN-RNA panel, conserved regions of gene segments that were specific to each targeted virus were identified from alignments of nucleotide sequences of strains available from the GenBank nucleotide sequence database (https://www.ncbi.nlm.nih.gov/nucleotide/). Alignments of multiple sequences were performed using Clustal Omega on Geneious Version 2022.2.2. or MUSCLE v5 [95]. Accuracy of the alignment results was confirmed by querying sequences using Basic Local Alignment Search Tool (BLAST, https://blast.ncbi.nlm.nih.gov/Blast.cgi, [96]) and comparing the query sequences and results. The accuracy of consensus sequences in finding the desired agent was examined on BLAST. Consensus sequences of conserved gene regions with pairwise identity percentages of 90% and above, and identical site coverage of 85% and above were selected. Consensus sequences that met the requirements from the last step were used as query sequences to search for similar biological sequences in GenBank using BLASTN. Qualified consensus sequences that had more than 50% non-specific similarities, those with query sequence coverage being less than 100%, and those with queried sequences’ identity percentage of less than 98% were removed. For the TICKHUNTER panel, the genes or genomic regions targeted for pathogen identification and subtyping were chosen based on highly conserved genes or genomic regions, genes used for speciation and MLST, and genes related to pathogenicity in certain species/genera [43, 88, 93, 97–104] (Suppl. Table 6).

#### Panel design and synthesis

The TITAN-RNA and TICKHUNTER panels were designed and synthesized by Twist Biosciences (South San Francisco, California). Baits in each panel consist of 120 bp biotinylated DNA segments based on the curated target sequences. For validation purposes, each panel was split into different versions. TITAN-RNA v1 (Suppl. Table 2) primarily targets known domestic and endemic TBVs, and TITAN-RNA v2 (Suppl. Table 3) primarily targets foreign tick-borne viruses. TITAN-RNA v3 is a final panel designed for syndromic diagnosis of fevers of unknown origin. It combines the targets from TITAN-RNA v1 and v2 with additional viruses linked to vector-borne febrile illnesses (Suppl. Table 4). TICKHUNTER v1 targets 26 genes or genomic regions of 19 tick-borne bacteria and three endoparasites (Suppl. Table 5). Highly conserved genes or genomic regions, as well as genes related to pathogenicity, like the surface antigen A of *Bab. microti* [105], were included. TICKHUNTER v2 includes all the targets from v1 plus additional genes used for the MLST of *A. phagocytophilum* [43] and *Bo. burgdorferi* [106] enabling association with known pathogenic lineages. In this panel, 55 genes or genomic regions of 19 bacteria and 5 endoparasites are targeted (Supp. Table 6)

### Library preparation and hybridization capture

Hybridization capture consists of three main steps: 1. DNA library preparation, 2. Hybridization capture (HC), and 3. NGS using the Illumina platform (See Suppl. Table 1 for a detailed protocol). For library preparation, the Twist Library Preparation Enzymatic Fragmentation Kit (1, 2.0) and the Twist Unique Dual Index (UDI) were used following the standard protocol, starting with 50 ng of DNA. These UDIs are compatible with the Illumina platform. For the HC, the Twist Standard Hybridization and Wash Kit v2, together with any of the designed panels, was used following the standard protocol, with a hybridization time ranging from 15 to 17 hours. After HC, the obtained enriched libraries were sequenced in the MiSeq (a medium-pass sequencer) or the iSeq (a low-pass sequencer).

### Analytical method validation

#### Design and synthesis of synthetic dsDNA

To assess the linearity and limit of detection (LOD) of each panel, synthetic dsDNA containing the known sequences of 11 genes or gene segments in total included in the panels was used. For the TICKHUNTER v1 panel, 8 gene targets from eight different TBPs, including *Rickettsia* spp. (or Panrickettsia), *Rickettsia rickettsii, Bo. burgdorferi, Bo. miyamotoi, Ehrlichia chaffeensis, A. marginale, A. phagocytophilum, and Bab. microti* were included (Suppl. Table 5). For the TITAN-RNA v1 panel, 3 gene targets from *Heartland virus* (HRTV), *Dabie bandavirus* (DBV), and POWV were included (Supplemental Data 9). DNA targets of 629 base pairs (bp) and 986 bp in length was synthesized for the 8 targets of the TICKHUNTER v1 panel and the three targets of the TITAN-RNA panel, respectively, by Integrated DNA Technologies (IDT, Newark, NJ, USA). Two pools of synthetic dsDNA were prepared, one containing the eight bacterial and parasite genes and the other containing the three viral genes. Both pools had an initial concentration of 10^5^ DNA copies/ μL of each synthetic dsDNA included in the pool. The initial concentration of each pool was confirmed with digital droplet PCR (ddPCR) using the ddPCR Supermix for Probes or the 1-Step RT-ddPCR Advanced Kit for Probes (BioRad; Hercules, California, USA) using the primers and probe listed in Suppl. Table 17. The manufacturer’s standard methods were followed for the ddPCR using the Droplet Generator Oil for Probes (BioRad), the QX200 Droplet Generator (BioRad), and the QX200 Droplet Reader (BioRad). DNA concentration was measured using the QX Manager software (BioRad).

#### Assessment of linearity and limit of detection of the panels

To assess the linearity of TICKHUNTER v1 panel, 10-fold serial dilutions from 10^5^ to 10^-1^ copies/μL were prepared from the pooled synthetic dsDNA containing eight selected genes. Seven replicates of each dilution series from 10^5^ to 10^-1^ copies/μL were performed. To assess the linearity of detection of the TITAN-RNA v1 panel, the Quantitative Synthetic POWV lineage II RNA (ATCC, VR-3275D) was used in 10-fold serial dilutions from 1×10^2^ copies per microliter to 1×10^-4^ copies per μL. Two replicates for each RNA dilution series were performed.

Limit of detection (LOD) testing was performed to compare the low-level detection capabilities of each panel. For the LOD assay of the TICKHUNTER v1 panel, 2-fold serial dilutions using nuclease-free water from 5 to 0.078125 DNA copies/μL of the eight pooled genes were performed five separate times. For the LOD assay of the TITAN-RNA v1 panel, the dsDNA of POWV NS5, HTRV L, or DBV L was serially diluted 1:2 from 1.7×10^-2^ copies/µl to 2.65625×10-5 copies/µl in nuclease-free water in six replicates per dilution. In addition, the Quantitative Synthetic POWV lineage II RNA (ATCC VR-3275SD) was serially diluted 1:2 from 0.225 copies/µl to 3.6×10^-3^ copies/µl in nuclease free water in six replicates per dilution.

To compare the LOD of the TITAN-RNA v1 panel and that of qPCR, synthetic POWV lineage II RNA was serially diluted 1:2 from 5 copies/µl to 1.95×10^-4^ copies/µl in nuclease free water in at least 5 repicates per dilution. POWV NS5 DNA was serially diluted 1:2 from 50 copies/µl to 3.125×10-5 copies/µl in at least five replicates per dilution. The qRT-PCR was done using the TaqMan Fast Virus 1-Step Master Mix for qPCR (Thermo Fischer Scientific, cat# 4444432). Four µl of each genomic DNA or RNA dilution was mixed with 9 µl nuclease-free water, 5 µl TaqMan Fast Virus 1-step Master Mix (4X), forward primers (10 µM), reverse primer (10 µM), and probe (10 µM). qRT-PCR was run on the QuantStudio™ platform (Thermo Fisher Scientific). Amplification thresholds were set at 10% of highest amplification in each run. Primer sequences in Suppl. Table 17.

To better approximate the complexity of clinical samples, an LOD experiment for each panel was performed using quantitative DNA or RNA standards at known concentrations spiked into blood and ear skin biopsy samples from specific-pathogen-free laboratory mice (FVB/NJ, JAX #001800). These mice were being euthanized for an unrelated study approved by the Cornell University Institutional Animal Care and Use Committee (IACUC, Protocol 2007-0115). For the TICKHUNTER v1 panel, the genomic DNA standard for *Bab. microti* (ATCC PRA-398DQ) was used. For the TITAN-RNA v1, the Quantitative Synthetic POWV lineage II RNA (ATCC, VR-3275D) was used. Mouse blood and ear skin biopsy samples were spiked with the genomic DNA standard for *Bab. microti* at final concentrations in each sample spanning two-fold serial dilutions from 10 to 0.078125 DNA copies/μL. Quantitative Synthetic POWV lineage II RNA was spiked serially 1:2 into mouse blood at 0.225, 0.1125, 0.05625, 0.028125, 0.0140625, 0.00703125, and 0.003515625 copies/μL, and into mouse ear skin biopsy at 0.45, 0.225, 0.1125, 0.05625, 0.028125, 0.0140625, 0.00703125, and 0.003515625 copies/μL. Isolation of DNA/RNA was performed using the MagMAX CORE Nucleic Acid Purification Kit (Applied Biosystems; Waltham, MA, USA) for blood samples, or the Quick DNA/RNA Pathogens MagBead Kit (Zymo Research) for the ear skin biopsy samples. Both extractions were performed using the KingFisher Apex (Thermo Fischer Scientific), following the suggested standard method for each kit.

The ratios of positive to negative samples were used to determine the fraction-positive value for each replicate and plotted using ggplot2 v.3.5.2 [107] in RStudio v.4.4.3 [108] or Quodata PROLab POD (Dresden, Saxony, Germany) [109]. A Sigmoidal Fitting model was fit with the x-axis as Log10 of POWV total copies and the y-axis as fraction-positive samples. The LOD was calculated as the concentration at which the fraction-positive samples decreased below 95%. A complementary log-log regression model for skewness was used to extrapolate an integer LOD where probit regression modelling was limited [41, 42].

#### Assessment of sequence variation tolerance of the TITAN-RNA panel

To assess the effects of natural and artificial variation in targeted genes on detection and sequencing using the TITAN-RNA v1 panel, synthetic dsDNA with varying degrees of divergence from a reference sequence was designed. POWV NS5 (NCBI # OL704303.1) was used as a baseline (0%) to design modified sequences that were 10%, 20%, 30%, 40%, and 50% variable using a custom Python code. A total of sixteen sequences, each 2,709 nucleotides long, were designed: five (10%), four (20%), three (30%), two (40%), and two (50%) independent replicates of randomly varied sequences (Supplemental Data 9). To ensure that the sequences were sufficiently divergent from the baseline reference sequence, code was written to determine the required number of mutations based on the original sequence length. POWV NS5 is 2709 nucleotides long, so a 10% different sequence will have approximately 271 mutations. The code was written to create mutations that were random, evenly distributed, and with no more than 10% overlap between variable sequences. The python code used to generate the sequence can be found in the data repository. The percent dissimilarity between the baseline and variant NS5 sequences is shown in Extended Data Fig. 3. To test how naturally divergent sequences will influence detection and sequencing using the TITAN-RNA panel, two sequences from each variant percentage group were baited with the NS5 gene sequences of genetically distant TBEV (NCBI# NC_001672.1) and ZIKV (NCBI# NC_012532.1) were also included. The artificially designed sequences and the naturally divergent sequences were ordered as high-fidelity dsDNA from IDT (2,709 bp). Each lyophilized sample was resuspended in nuclease-free water at 10 ng/μL in nuclease-free water. The final concentration of each sequence stock was measured using a Qubit Fluorometer High Sensitivity Kit (Thermo Fisher Scientific), and total copies per microliter were calculated based on the sequence length. Ten thousand total copies of each sequence were spiked into individual aliquots of 40 μL water with 50 ng of clean mouse ear skin biopsy DNA. Hybrid-capture was performed on up to nine replicates and analyzed as described. The average coverage depth was calculated as follows:

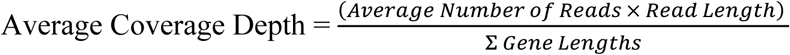

### Field sampling

To test the diagnostic performance of the panels for pathogen detection in clinical samples, blood and skin biopsy (ear notch) samples were collected from white-footed mice and raccoons in Queens and Staten Island, two boroughs of New York City (NYC) known to have low (Queens) and high (Staten Island) TBP circulation, with multiple vector species established [110] (Suppl. Table 10). Field sampling was approved by the IACUC of Columbia University (AC-AABM9557 and AC-AABS3611), Cornell University (2022-0198), and the US Army Animal Care and Use Review Office (ACURO #TB210019.e001). A License to Collect was also obtained from the New York State Department of Environmental Conservation (License # 2704).

White-footed mice and raccoons were live-trapped and sampled from early August to early September 2023. This time period corresponds with when larval *I. scapularis, H. longicornis,* and *A. americanum* ticks actively search for a host blood meal in NYC [111]. For all individual white-footed mice and raccoons captured, all parasitizing ticks were collected, a small (2mm) ear tissue biopsy was taken, blood was collected, and all animals were tagged using metal ear tags before being released back to the trapping location. Tick and tissue samples were placed into prefilled vials with RNA/DNA Shield (Zymo Research, Irvine, CA) and stored at 4°C until further processing. White-footed mouse blood was collected and stored in EDTA microtubes at-80°C, and raccoon blood was collected and stored in DNA/RNA Shield blood collection tubes and stored at 4°C until further processing. All ticks collected from white-footed mice and raccoons were identified to species and life stage using microscopy [112, 113].

Host-seeking ticks were collected and identified in June 2022, 2023, and 2024 as part of the New York State Tick Blitz citizen science project, as previously described [50]. Briefly, ticks were collected one time per day to reduce the likelihood of transporting ticks to other locations, and they were collected along a 300-m linear transect, either continuous or divided into shorter parcels in multiple sites across New York State. Nymphal and adult ticks were collected from cloth drags with forceps and placed in plastic vials. Larvae were collected with lint rollers and then placed into plastics bags. Ticks were refrigerated in plastic sealable bags with moistened paper towels. Live ticks were mailed to Cornell University for identification as previously described [50].

Sera was collected from meso-mammals (144) and small-mammals (44) in parks located in Queens, Staten Island, and Brooklyn, totaling 188 samples. Specifically, five opossums, 11 skunks, 44 mice, and 128 raccoons were sampled. All samples were stored at-80°C until processed.

#### Nucleic acid extraction from field samples

In total, 64 ear skin biopsies and 39 blood samples from white-footed mice, and 24 blood samples from raccoons were processed (Suppl. Table 10). DNA and RNA were co-extracted from ear skin samples using the Quick-DNA/RNA Pathogens MagBead Kit (Zymo Research) as mentioned above for the ear notches of control mouse samples. DNA from mouse blood was extracted using the Monarch Genomic DNA Purification kit (New England Biolabs, NEB), using the standard protocol for whole blood extraction. DNA and RNA from raccoon blood samples were extracted using the MagMAX CORE Nucleic Acid Purification Kit (Applied Biosystems) following the whole blood workflow. Prior to extraction, blood samples were equilibrated to room temperature for one hour. The extraction process was performed on the KingFisher Apex (Thermo Fischer Scientific), following the standard method. The DNA extracted from the blood and ear notch samples was used for qPCR and library preparation, followed by HCNGS.

Questing ticks collected between 2022 and 2024 in New York State were homogenized using a 2.0-mm hollow brass bead in 800 uL of PBS and the BeadBlaster D2400 (Benchmark Scientific; Sayreville, NJ, USA) at 2.34 meters per second for 5 minutes. Homogenized samples were then centrifuged at 14,000 rpm for 1 minute and incubated for 1 hour with Proteinase K. Samples were centrifuged again for 3 minutes at 14,000 rpm, and 300 uL supernatant was used for the subsequent DNA/RNA extraction on the KingFisher Apex (Thermo Fisher Scientific) using the Quick-DNA/RNA Pathogen MagBead extraction kit (Zymo Research, catalog number # R2145 or R2146). Nucleic acid samples eluted from tick homogenates were frozen at-80°C.

#### qPCR for pathogen detection from field samples

The DNA extracted from raccoon, white-footed mouse, and tick samples was used to perform gold standard qPCR assays to detect *Bo. burgdorferi, Bab. microti, A. phagocytophilum,* and *Bartonella* spp [103, 114–116]. The real-time PCR was performed on the Applied Biosystems QuantStudio 3 Real-Time PCR System (Applied Biosystems, USA). Each reaction contained TaqMan™ Fast Virus 1-Step Multiplex Master Mix (Thermo Fisher Scientific, USA), the primers and probes listed in Suppl. Table 17. Reactions were performed using the following thermocycling conditions: 50°C for 2 min, 95°C for 10 min, followed by 45 cycles of 95°C for 15 s and 60°C for 1 min. For POWV, qRT-PCR was done using the TaqMan Fast Virus 1-Step Master Mix as described for LOD. *Diagnostic accuracy estimation.* Sensitivity and specificity of real-time PCR and HCNGS for *Borrelia* were estimated by Bayesian latent class analysis under the Hui-Walter paradigm, which identifies both parameters from the cross-classified results of two imperfect tests applied to two or more populations of differing prevalence, without assuming a reference standard [117]. Ear skin biopsies from 64 white-footed mice were partitioned by borough (Queens, n=39; Staten Island, n=25). The model estimated six parameters (sensitivity and specificity of each assay, and prevalence in each population) from six degrees of freedom, and is therefore identified. Tests were assumed conditionally independent given true infection status, and sensitivity and specificity were assumed constant across populations. A non-informative Beta(1,1) prior was used for sensitivity and specificity of both tests. Prevalence priors were determined as a most-likely value (mode) together with a one-sided confidence bound and converted to Beta distributions using the epi.betabuster() function in the package epiR, R version 4.3.2 [118]. Queens prevalence was elicited as most likely 15% with 95% certainty below 45%, and Staten Island as most likely 45% with 95% certainty below 75% based on literature estimates [119, 120]. This resulted in a Beta (α=2.235, β=7.998) prior distribution for prevalence in Queens and a Beta (α=3.497, β=4.052) prior distribution for prevalence in Staten Island. The latent class model was run in R using JAGS [121] through the R2jags package and outputs visualized using the packages Coda and MCMCvis. To investigate the conditional independence assumption between tests because since both assays target pathogen DNA, the analysis was also conducted using a LCA model that accounts for dependence between tests. However, since the credible interval for positive covariance and negative covariance both spanned 0, there was no strong evidence for conditional dependence between tests and the independent model was used.

#### Hybrid capture sequencing workflow

RNA was converted to cDNA using SuperScript IV VILO Master Mix with ezDNase (Thermo Fischer Scientific, cat# 11766050) and converted to dsDNA using DNA Polymerase I, Large (Klenow) (NEB, Ipswich, MA, USA; cat# M0210L). Libraries were prepared using the TWIST Bioscience Library Preparation EF 2.0 with Enzymatic Fragmentation and the Illumina TruSeq-compatible Twist CD Index Adapter Set 1-96 (Twist Biosciences). Per the standard protocol (Extended Data Fig. 1), the target DNA concentration for each library preparation was set to less than or equal to 50 ng in 40 µL of nuclease-free water, and the DNA fragment target was approximately 300 bp. The final concentrations of the libraries were quantified with Qubit 1X dsDNA using Broad Range or High Sensitivity assay kits (Invitrogen; Waltham, MA, USA), and DNA concentrations measured using the Qubit Flex Fluorometer. Eight libraries were pooled at a time for multiplexing to prepare for hybrid-capture. DNA in pooled libraries was ‘captured’ according to the Twist Target Enrichment Standard Hybridization v2 protocol. The custom bait panel was used to hybridize each pool for 15 to 17 hours and sequenced on the MiSeq Sequencing System (Illumina) using the MiSeq Reagent Kit v3 (Illumina) and a read length of 151 or 251 base pairs. After sequencing, the raw reads were initially screened with the Chan Zuckerberg ID (CZID) cloud-based platform v3.0.2 (https://czid.org/; [122]). To verify the CZID positives, raw reads were trimmed with Trimmomatic v.0.39 [123] and mapped to the reference sequences using BWA-MEM2 v.2.2.1 [124, 125], on the Galaxy cloud informatics platform [126]. Reference-free assemblies were built to confirm the closest sequence match.

#### MLST, phylogenetic analyses, and sequencing platform comparison

For *A. phagocytophilum* and *Bo. burgdorferi,* the complete or partial sequences obtained from the genes targeted for MLST were uploaded to the *A. phagocytophilum* database [43] or the *Bo. burgorferi* database [106] of the Public Databases for Molecular Typing and Microbial Genome Diversity (PubMLST [127]), respectively. Each *A. phagocytophilum* and *Bo. burgdorferi* detected was typed following their respective MLST scheme to generate an allelic profile [43, 106]. This profile was used to find the allelic groups (also known as sequence types or STs) that are identical or closely match those of the samples from this study. The partial or complete sequences of the seven genes of each *A. phagocytophilum* obtained in this study were concatenated and aligned with those of the concatenated sequences of the closely matched STs using the Clustal Omega algorithm in Geneious [106]. This alignment was used to construct a maximum likelihood (ML) phylogenetic tree in MEGA11 [128]. For the samples in which the sequence of the 23S rRNA of *Babesia* spp. and the *ssrA* gene of *Bartonella* spp. were obtained, a BLAST search was performed to find closely related sequences. For *Bo. burgorferi,* the three complete sequences of the *ospC* gene obtained in this study were aligned with the sequences of the 33 major groups (MG) of *Bo. burgdorferi* sensu stricto [129]. For these three pathogens, the sequences were aligned, and an ML phylogenetic tree was constructed in the same manner as for sequences of *A. phagocytophilum*.

### Divergent sequence tolerance

To measure the sequence variability tolerance of the TITAN-RNA v1 panel system and conserve resources, nine of the sixteen randomly altered NS5 “variant” sequences described in 2.1 (10seq1, 10seq2, 20seq1, 20seq2, 30seq1, 30seq2, 40seq1, 50seq1, 50seq2), and NS5 of TBEV (NCBI# NC_001672.1) and ZIKV (NCBI# NC_012532.1), were resuspended to a concentration of 10 ng/μL in nuclease-free water. The final concentration of each sequence stock was measured using a Qubit Fluorometer High Sensitivity Kit, and total copies per microliter were calculated based on the sequence length. Ten thousand total copies of each sequence were spiked into individual aliquots of 40 μL water with 50 ng of clean mouse ear skin biopsy DNA. Hybrid-capture was performed and analyzed as described. The average coverage depth was calculated as follows:

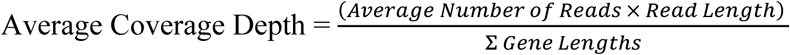

**Virus discovery.** *H. longicornis* samples were screened with both untargeted and targeted methods. Untargeted deep RNA sequencing was performed in separate laboratory (Cornell University Transcriptional Regulation and Expression Facility; TREx RRID: SCR_022532). After DNAse treatment and rRNA depletion of 20ng total RNA with the QIAseq FastSelect-rRNA HMR and 5S/16S/23S kits (Qiagen; Germantown, Maryland, USA), RNA sequencing libraries were generated using the NEBNext Ultra II Directional Library Prep kit (New England Biolabs, Ipswich, MA, USA), quality checked, and 100M reads were targeted per sample using 2 × 150 bp paired-end sequencing on the Illumina NovaSeq 6000 platform with the S4 flow cell. Raw reads were trimmed, assembled, and mapped to references from the TITAN-RNA v1 panel using GalaxyTrakr’s Trimmomatic, rnaviralSPADES v3.15.5 [130–132], and BWA-MEM2 tools. Contigs were also screened for other viral homologies using VirBot [133].

Conventional RT-PCR or nested-RT-PCR and long-read amplicon sequencing was used to further characterize viral sequences. RT-PCR or nested-RT-PCR was performed with the SuperScript III or IV One-Step RT-PCR System (Invitrogen) with FLSV-US primers and probe listed in Supplemental table 18. Ten microliters of PCR product were run on 1% agarose gels, imaged with a ChemiDoc MP Imaging System (BioRad). Gels were extracted with QIAquick Gel Extraction Kit (Qiagen, catalog# 28704), or RT-PCR products were directly cleaned with QIAquick PCR Purification Kit (Qiagen, catalog# 28104). PCR products were quantified using Qubit 1X dsDNA High Sensitivity assay kits and concentrations measured by fluorimetry (Qubit Flex). Amplicons were barcoded using the Native Barcoding Kit 96 V14 (Oxford Nanopore; Oxford, UK; cat# SQK-NBD114.96I), prepared libraries were loaded onto a MinION R10.4.1 (FLO-MIN114) flow cells and sequenced with a MinION Mk1b Oxford Nanopore Portable Sequencing device. Raw reads were trimmed, assembled, and mapped to references from the TITAN-RNA v1 panel using GalaxyTrakr’s Porechop v.0.2.4 [134], Flye v.2.9.5 [135, 136], and BWA-MEM2 tools, respectively. Sanger sequencing performed by Cornell Biotechnology Resource Center. Hybrid assemblies from TITAN-RNA, Sanger, and Nanopore-based sequencing reads were built with Geneious Prime de novo assembler. Sequences were analyzed with Prokka v.1.14.6 [137, 138], and maximum likelihood phylogenetic trees made using GalaxyTrakr’s IQtree v.2.4.0 [139–145], and visualized with iTOL v.7.2 [146].

The codon adaptation index (CAI) was determined to evaluate the relative adaptability of the codon usage of a virus gene towards that of highly expressed genes within a given host [47]. The maximum CAI is 1 and the minimum is 0. In general, the higher the CAI value, the more efficient the mRNA can be translated. The CAI values of the glycoprotein, envelope and NS5 genes from the novel segmented *Flavi-*like virus (FLSV-NY) were analyzed in the context of the following: *Homo sapiens* as host (NCBI accession no. GCF_000001405.40); several known or potential reservoirs hosts species implicated in the transmission: *Canis lupus familiaris* (dog, GCF_011100685.1), *Peromyscus leucopus* (white-footed mouse, GCA_004664715.2), *Peromyscus maniculatus bairdii* (prairie deer mouse, GCF_049852395.1), *Procyon lotor* (racoon, downloaded from https://www.dnazoo.org/assemblies/procyon_lotor) [147]; the main vector responsible for the transmission in sylvatic and urban cycle is *Haemaphysalis longicornis* tick (Asian longhorned tick, GCA_013339765.2) and other tick vectors to test for expanded range or transmission potential: *Amblyomma americanum* (lone star tick, GCA_030143305.2), *Amblyomma maculatum* (Gulf Coast Tick, GCA_023969395.1; coding sequences from [148]); *Dermacentor andersoni* (Rocky Mountain Wood tick, GCF_023375885.2), *Dermacentor variabilis* (American dog tick, GCF_050947875.1), *Ixodes pacificus* (Western blacklegged tick, GCA_964199305.2), *Ixodes scapularis* (Eastern blacklegged tick, GCF_016920785.2), *Rhipicephalus sanguineus* (brown dog tick, GCF_013339695.2). As reference, we also tested the glycoproteins VP1a and VP1b genes of ALSV, CJLV and JMTV, as well as the NS5 gene of POWV, TBEV, and YFV (Supplemental table 19). To be noted, although accession number MN095520.1 isolate is classified as JMTV, its glycoproteins show much higher similarity to ALSV (>99% by BLAST), and was therefore included in the latter when summarizing CAI statistics. Because eukaryotic viruses generally do not have their own translation machinery, they typically evolve codon usage in response to their host tRNA pools, thus viruses evolved their increased translation efficiency match the most abundant cognate tRNA. Gene expression is proportional to respective tRNA anticodon genome copy numbers [149–153]. Thus, we have identified the highly expressed genes for each host were inferred by calculating the tRNA adaptation index (tAI) [149–153] of host CDS sequences at least 500 base pairs long: sequences with no start codon or having an internal stop codon were excluded; these filters were implemented in order to reduce the chance of pseudogenes being part of the data set. tAI were considered highly expressed from the top 500 genes [154] excluding single triplet amino acids (Met and Trp) to reduce bias analyses in favor of proteins rich in such amino acids, as they would get a high CAI [155]. Afterwards, we calculated the CAI for each target viral sequence. Analyses were performed in R v.4.4 and RStudio. Fasta files were imported into R using the Biostring package [156]; tAI and CAI were calculated using the R package cubar [157] and CAI plots were drawn with the ggplot2 package [107]. Because of missing tRNA annotations, tRNA anticodons for *A. americanum*, *A. maculatum*, *H. longicornis*, *I. pacificus*, and *P. lotor* were inferred using tRNAscan-SE v.2.0.12 [158]. Supplemental Codon Adaptation Index was calculated using the E-CAI server (https://ppuigbo.me/programs/CAIcal/E-CAI/; [48]) using the *Homo sapiens* codon usage table (https://www.kazusa.or.jp/codon/cgi-bin/showcodon.cgi?species=9606).

### Bloodmeal analysis

Amplification of tick-associated host 16S mitochondrial rRNA was performed as described in our previous work [159], using the following untagged primers: UnivMamF: 5’-GACGAGAAGACCCTATGGAGC-3’, UnivMamR: 5’-TCCGAGGTCGCCCCAACC-3 [160]. In brief, target regions (98 to 120 bp) were amplified using Platinum II Hot-Start Green PCR Master Mix (Invitrogen, Carlsbad, California), and samples were visualized in 2% Agarose gel. Samples with bands between 98-120bp in size were purified with a 1.8x bead to sample ratio using Ampure XP beads (Beckman Coulter, Brea, California) and DNA libraries were prepared for sequencing on a MiSeq Sequencing System (Illumina). The raw paired-end demultiplexed FASTQ files were quality checked with FastQC and summarized with multiQC, then imported into QIIME 2 amplicon v2024.10 [161–163] via a PairedEndFastqManifestPhred33v2 manifest, to create a SampleData[PairedEndSequenceWithQuality] artifact. Quality and sequencing depth were summarized using ‘qiime demux summarize’. Adapters, primers, and low-quality bases were removed using qiime cutadapt (error-rate 0.1, p-times 1, p-overlap 3, p-minimum-length, p-quality-cutoff-5end/3end 0, p-quality-base 33) [123, 164] and again visualized with ‘qiime demux summarize’. Trimmed forward and reverse reads were merged using ‘qiime vsearch merge-pairs’ (p-minmergelen 60, p-maxmergelen 100), then dereplicated using ‘qiime vsearch dereplicate-sequences’, producing an amplicon sequence variant (ASV) feature table and representative sequences. Chimeric sequences were identified by first clustering like sequences using ‘qiime vsearch cluster-features-de-novo’ (p-perc-identity 0.99) followed by ‘qiime vsearch uchime-denovo’ to detect and remove chimeric sequences. A custom naïve Bayes classifier, trained on a curated mitochondrial 16S reference database trimmed to the amplicon region, assigned taxonomy to ASVs with ‘qiime feature-classifier classify-hybrid-vsearch-sklearn’ (confidence = 0.8). The resulting per-feature taxonomy was visualized with ‘feature-table summarize’ and ‘taxa barplot’ functions. Reads mapping to a maximum depth of the Chordata phylum were discarded, and those mapping to primates were removed as likely contaminants. Remaining reads were summed up and used to calculate relative abundance per blood meal taxa. To build the curated mitochondrial reference database for species that may have been a tick bloodmeal, the complete mitochondrial genomes for mammals, birds, reptiles, and amphibians were collected from NCBI RefSeq using ‘qiime rescript get-ncbi-data’ and an ID list pulled from EDirect queries such as’Mammalia[Organism] AND mitochondrion[Filter] AND refseq[filter] AND complete genome’ for each phylogenetic class. The mitochondrial genomes were then trimmed to the Universal Mammalian Primer amplicon region (∼75 bp) using ‘feature-classifier extract-reads’ (identity = 0.8) [159]. Separate classifiers were trained on representative sequences for each animal classusing the naïve Bayes learning classification algorithm ‘feature-classifier fit-classifier-naive-bayes’. Mammal, bird, reptile, and amphibian representative sequences were merged with ‘merge-seqs’, and sequence taxonomy was merged with ‘merge-taxa’. The merged sequences and taxonomy were used to train a naïve Bayes classifier with the same steps described to produce the individual classifiers. Classifiers were validated by applying ‘classify-sklearn’ to both representative sequences and sequences from blood of known animal sources, then inspecting confidence scores and read counts. Scripts for taxonomic classification from raw FASTQs and for building the taxonomic classifier can be found in the data repository. Additionally, *H. longicornis* sequences were extracted from raw RNAseq reads and mapped to a database of complete mitochondrial 16s rRNA sequences. Relative abundance was calculated as reads mapped to a given species over total reads mapped to the database.

### *Flavivirus* Seroprevalence mapping

The geographic distribution of *Flaviviridae* exposure in meso-, or small-mammals was monitored using antibodies detected in collected sera. Since there are currently no commercial screening kits available for any tick-borne *Flaviviridae* endemic to the US, the highly conserved nature of the *Flaviviridae* virion structure was exploited using a crude antigen FSME (TBEV) ELISA Kit (Arigo Biolaboratories, Taiwan, cat #ARG83078) [58–65]. ELISA data were first evaluated for normality using Shapiro-Wilk tests and homogeneity of variance using Bartlett’s tests. Data were visualized as box plots showing medians, interquartile ranges, and individual data points, with a color gradient from green (lowest) to red (highest), generated in RStudio v2025.05.1+513 using ggplot2 v3.5.2 and tidyverse v2.0.0. To reduce skewness a constant was added, and values were log-transformed using base R functions. Based on skewness observed in either variance and normality, Kruskal-Wallis, Dunn’s post-hoc tests, and effect size calculations were performed with *η²ₕ*using rstatix v0.7.3. Pairwise adjusted p-values (Padj) from Dunn’s tests were visualized as significance indicated by asterisks (*Padj < 0.05, **Padj < 0.01, ***Padj < 0.001). Summary statistics are available in Source data 8.

## Data Availability

Sequence data from this study are deposited in NCBI BioProjects PRJNA1271117 (TITAN-RNA) and PRJNA1270965 (TICKHUNTER). Additional data and code are available at the Cornell University Library eCommons Repository (DOI pending) and https://gitlab.com/goodmanlab.

## Extended Data Captions

1. **Extended Data Fig. 1**: Protocol overview
2. **Extended Data Fig. 2**: Bench validation

a. Linearity/dynamic range curves.
b. Limit of Detection.
3. **Extended Data Fig. 3**: Heatmap of random POWV variant stress test.
4. **Extended Data Fig. 4**: *Anaplasma* MLST phylogeny
5. **Extended Data Fig. 5**: Blood meal mitochondrial remnant analysis in host-seeking ticks

a. *Haemaphysalis longicornis mt16s*
b. *Haemaphysalis longicornis cytB*
c. *Ixodes scapularis mt16s*
d. *Ornithodoros hermsi mt16s*

## Acknowledgments

We are thankful to Dr. Lars Eisen and Dr. Joan Kenney for critical review of the manuscript. The authors also thank the de Mestre laboratory at Cornell University for assistance with ddPCR and quo data (https://www.quodata.de/) for use of their PCR analysis tool. Equipment used for this study was provided by the United States Food and Drug Administration’s Veterinary Laboratory Investigation and Response Network under Research Collaboration Agreement FY16-RCA-CVM-01-CU; the Transcriptional Regulation & Expression (TREx) Facility (RRID: SCR_022532) at the Cornell Institute of Biotechnology; and UF-ICBR Bioinformatics Core at the University of Florida (RRID:SCR_019120).

## Extended Data Figures

**Extended Data Figure 1.**
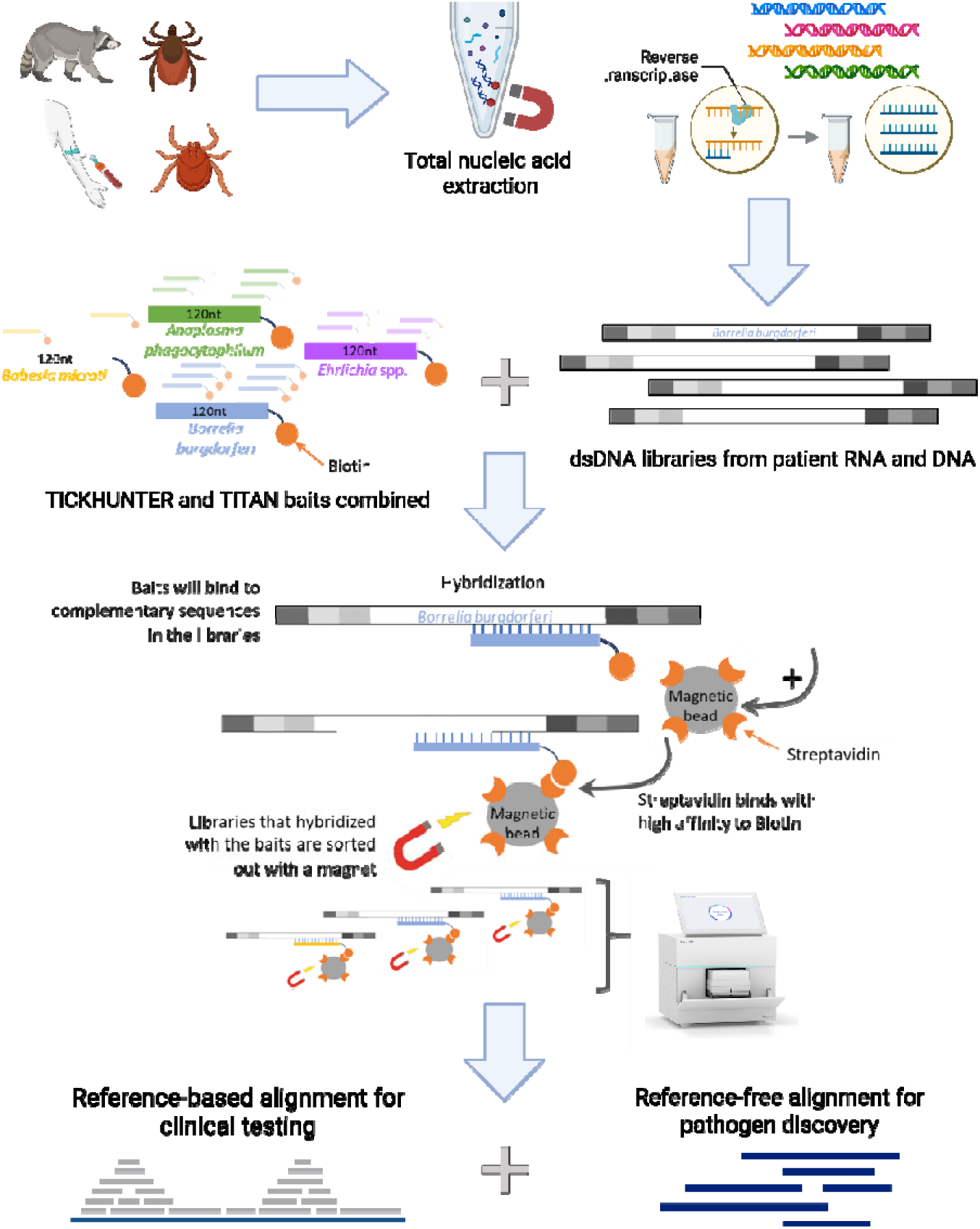
Overview of the combined hybridization capture next-generation sequencing method. This method comprises four main parts: library preparation from samples, a panel featuring biotinylated baits with specific target sequences, hybridization of complementary sequences between the library and the bait panel, followed by next-generation sequencing using Illumina platforms. This method can be used to detect and sequence known tick-borne pathogens (bacteria and parasites), including *A. phagocytophilum* (in green), *Ba. microti* (in yellow), *Bo. burgdorferi* (in blue), and *Ehrlichia* spp. (in violet).

**Extended Data Figure 2.**
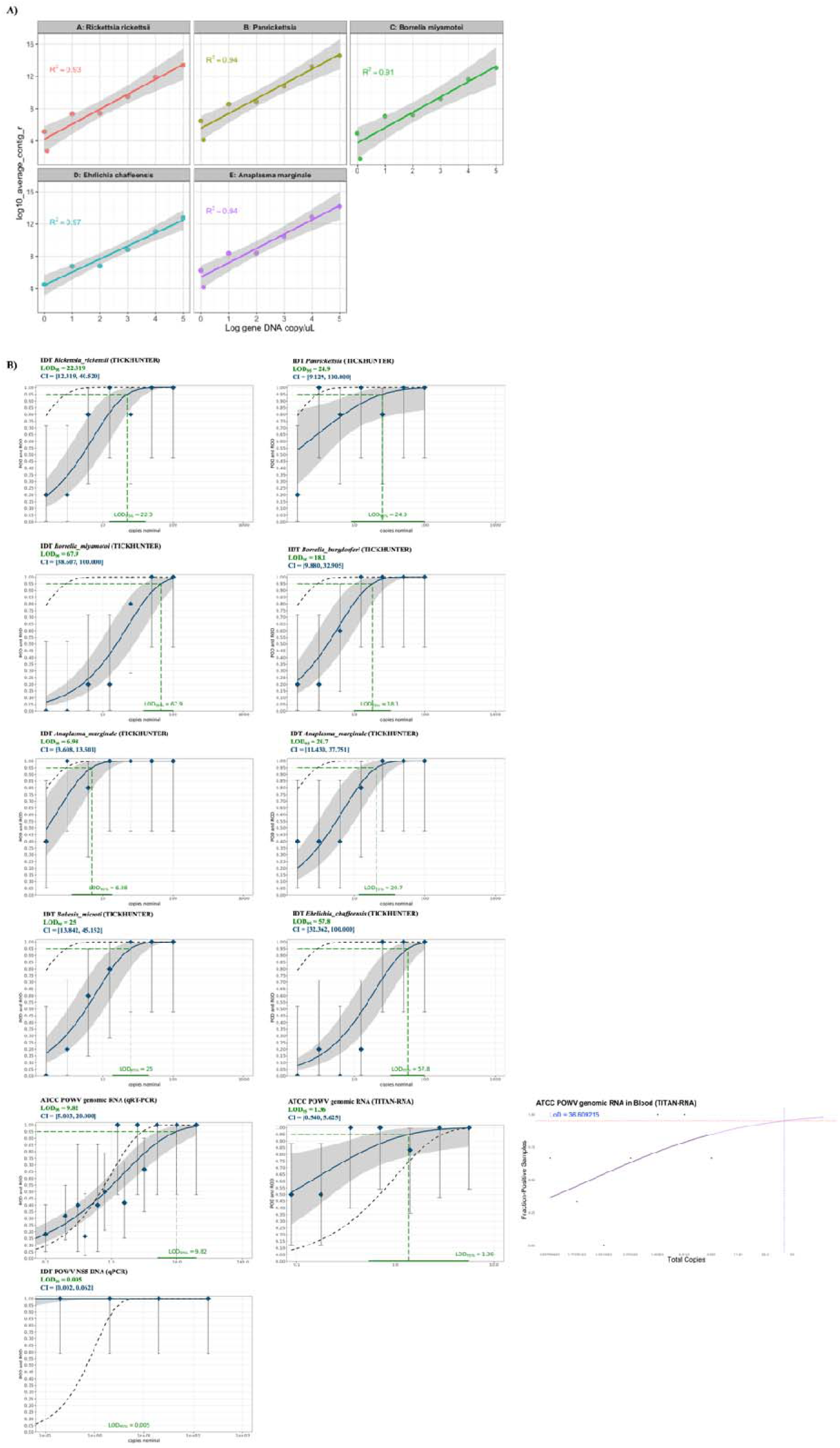
(A) Linearity curves for representative DNA or RNA pathogens plotted as Log_10_ of Mean Contig R (number of reads aligned per taxonomically matched contig, Y-axis) versus Log_10_ of Concentration (copies/µl, X-axis) (n ≥ 3). (B) Fitted sigmoidal curves plotted as fraction-positive samples versus total copies for the estimation of the LOD for each of the eight genes included in the pooled synthetic DNA or RNA (n = 3).

**Extended Data Figure 3.**
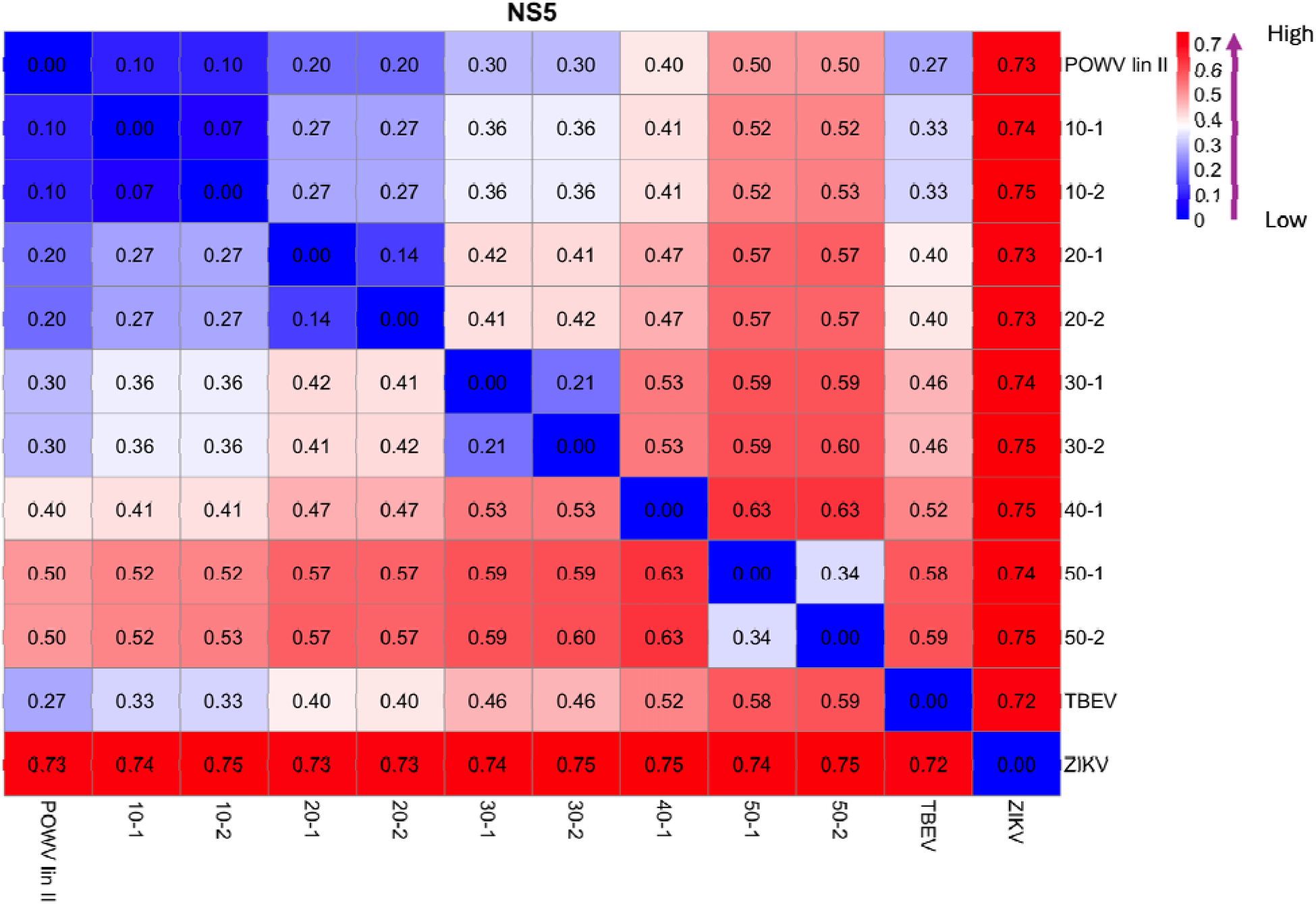
Heatmap of random POWV variant stress test

**Extended Data Figure 4.**
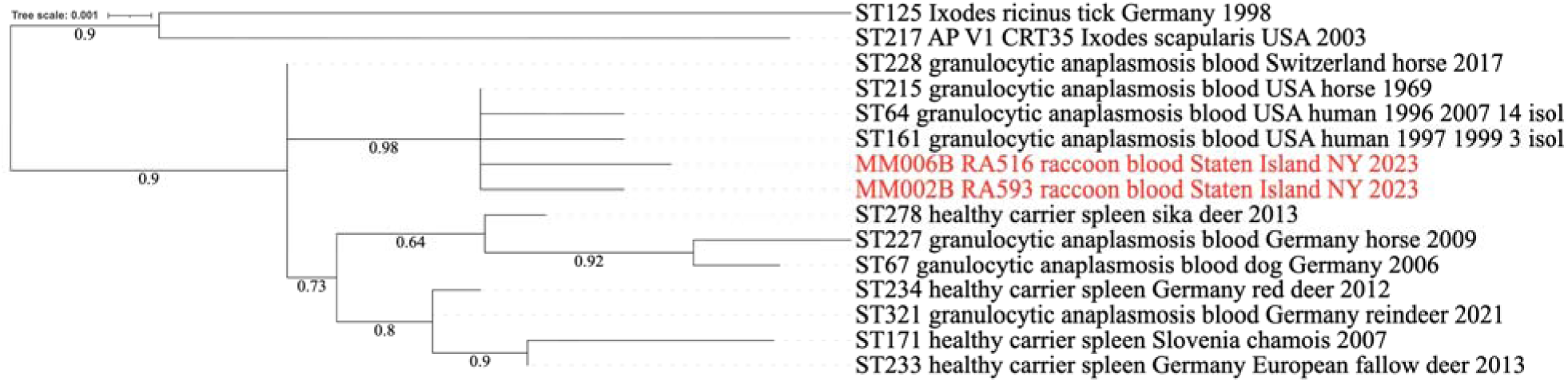
Maximum likelihood phylogenetic tree of five concatenated genes of the multi-locus sequence typing scheme of *A. phagocytophilum* (915 nt). Since it was not possible to obtain the complete sequence of the seven genes included in the MLST scheme for all the samples, we concatenated the sequences of five genes (*dnaN, glyA, mdh, pheS, sucA*) from two samples (RA516, RA593). These were the longest overlapping sequences obtained from these two samples, allowing for a more complete genetic comparison with other sequences. The two sequences obtained in this study are in bold and form a group with sequences from three allelic groups (ST215, ST64, ST161) detected in the USA in cases of granulocytic anaplasmosis from humans and a horse. An additional *A. phagocytophilum* detected in the USA exclusively in an *I. scapularis* tick is also shown (Ap-V1) and does not group with the sequences obtained in this study or those from human cases in the USA. The name of each sequence includes its allelic group number (ST), the health information of the host (granulocytic anaplasmosis or healthy carrier), the sample type (blood or tick), the host species, and the country and year of collection. If more than one isolate has been reported for a given ST, the number of isolates is indicated in parentheses. Numbers in the branches indicate bootstrap percentage values for 1,000 replicates. Branches with <50% support were collapsed. Nucleotide substitution model used: K2+G

**Extended Data Figure 5.**
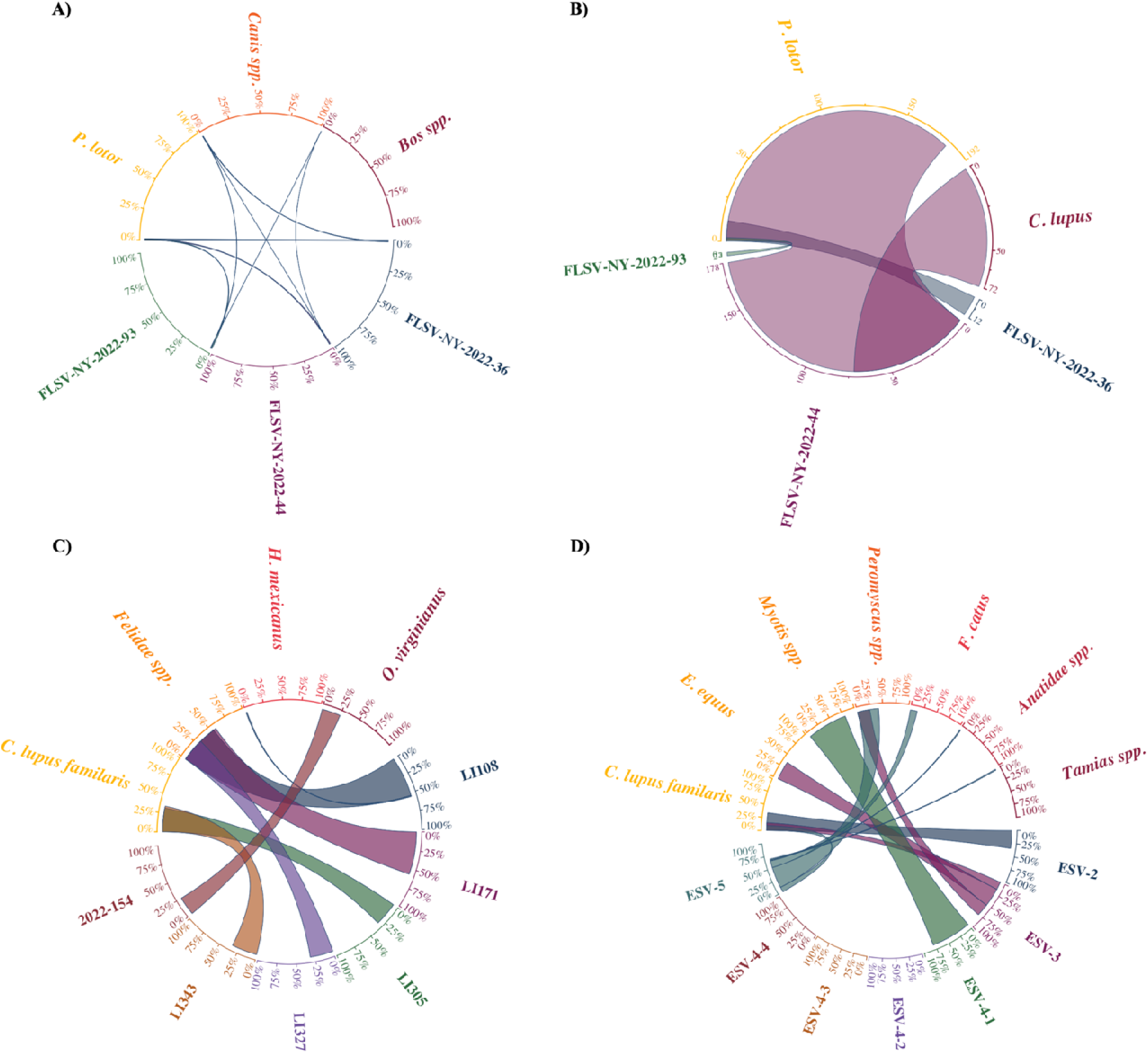
Blood meal mitochondrial remnants in host-seeking ticks (axes as relative abundance in percentage). A) RNA sequencing of FLSV-NY positive *Haemaphysalis longicornis* tick pools mapped to mitochondrial 16s rRNA database. B) RNA sequencing of FLSV-NY positive *Haemaphysalis longicornis* tick pools mapped to mitochondrial *cytB*. C) Mitochondrial 16s rRNA from POWV positive *Ixodes scapularis* based on metabarcoding. D) Mitochondrial 16s rRNA from *Ornithodoros hermsi* based on metabarcoding.

